# Spatially patterned cytoskeletal organization shapes astrocyte branch complexity

**DOI:** 10.1101/2025.10.10.681734

**Authors:** Meghan E. Wynne, William E. Barclay, Dharshini Gopal, Naznin Jahan, Johanna J. Bergstrom, Lana T. Ho, Eva A. Lopez, Eva Nogales, Meng-meng Fu

## Abstract

Astrocytes, one of the most abundant cell types in the brain, extend elaborate branches that enable diverse functions, from synapse maintenance to blood-brain-barrier integrity. The cytoskeletal basis of this architecture has remained unclear, since traditional culturing methods produce minimal branching. Using immunopanning and serum-free conditions, we generated primary rodent astrocytes with complex, hierarchically branched morphology and surveyed their cytoskeleton using confocal microscopy and cryogenic electron tomography. We show that microtubules in the primary branches of immunopanned astrocytes are oriented primarily plus-end-out. Proximally, microtubules appear stabilized by post-translational modifications (PTMs) and microtubule inner proteins. Distal regions lack stabilizing microtubule PTMs, and are enriched in intermediate filament GFAP. Additionally, diverse actin microstructures, including reticular webbing, extend astrocyte boundaries beyond the microtubule–GFAP framework. Finally, pharmacological disruption of actin polymerization alters primary branch number and length, providing functional support for the interplay of cytoskeletal classes in defining astrocyte branching. Together, our results uncover spatial principles of astrocyte cytoskeletal organization that support complex branching morphology.

## Introduction

Astrocytes make up ∼20–40% of cells in the mammalian brain (von Bartheld et al., 2016; Zhang et al., 2023). They perform essential roles in neurotransmission, synaptogenesis, synaptic pruning, maintenance of the blood-brain barrier (BBB), and inflammatory signaling (Fisher and Liddelow, 2024; Chung et al., 2024; Linnerbauer et al., 2020; Díaz-Castro et al., 2023). These wide-ranging functions rely on their structurally complex and highly branched morphology, which enables individual astrocytes to simultaneously interact with diverse cell types, such as neurons, vascular cells, oligodendrocytes, and microglia (Baldwin et al., 2024; Torres-Ceja and Olsen, 2022). Astrocyte morphology also dynamically changes in response to environmental cues (Schiweck et al., 2018; Lawal et al., 2022), transitioning from a highly-branched homeostatic state to a more amoeboid reactive state with hypertrophy of branches following injury or disease (Escartin et al., 2021; Schiweck et al., 2018; Sofroniew, 2020). Although the structural complexity of astrocytes underlies their functions, the cytoskeletal organization that drives the formation, maintenance, and plasticity of astrocyte branches remains obscure (Torres-Ceja and Olsen, 2022; Weigel et al., 2021).

A key unresolved question in astrocyte cell biology is how microtubule polarity is organized across astrocyte branches (Weigel et al., 2021). Microtubules, which serve as tracks for long-range transport of diverse cargos, have polar structures that define the directionality of transport by microtubule-based motors. Microtubule organization and polarity are critically important in cells with complex branches, where specialized cargo needs to be delivered to distal branches that can extend far from the soma (Cason and Holzbaur, 2022; Rolls, 2022; Burute and Kapitein, 2019; Barlan and Gelfand, 2017). In the 1980s and 1990s, researchers assessed microtubule polarity using the “hooking” technique, which visualizes the orientation of assembling tubulin subunits via electron microscopy (Baas and Lin, 2011). More recently, live imaging of microtubule plus-end binding (EB) proteins and cryoelectron tomography (cryo-ET) have largely replaced hooking. In neurons, hooking (Baas et al., 1988), live-cell imaging in culture (Yau et al., 2016; Stepanova et al., 2003), and *in vivo* imaging in rodents (Yau et al., 2016; Kleele et al., 2014) show that dendrites contain mixed-polarity microtubules (∼65–70% plus-end-out), while axonal microtubules are almost uniformly plus-end-out, a finding confirmed by cryo-ET (Bouchet-Marquis et al., 2007; Foster et al., 2022; Hoffmann et al., 2021). Like axons, microtubules in oligodendrocyte processes are also oriented with plus-end-out polarity (Lunn et al., 1997; Fu et al., 2019). Live imaging has also recently demonstrated that microtubule polarity in microglia, the brain’s resident immune cells, dynamically shifts with immune activation (Rosito et al., 2023; Adrian et al., 2023). Though microtubule polarity is defined for neurons and other glia, it has remained poorly defined for the structurally complex astrocyte. Determining microtubule polarity in morphologically complex purified astrocytes is therefore an important step toward understanding how cytoskeletal organization supports astrocyte branch architecture and compartmentalized function.

In addition to microtubules, astrocytes also express actin and intermediate filaments (IFs), including glial fibrillary acidic protein (GFAP), which is commonly used as an astrocyte marker and often becomes upregulated in reactive states (Sofroniew, 2009; Escartin et al., 2021; Messing and Brenner, 2020). However, few studies have explored the spatial organization and interplay between these three classes of the cytoskeleton in astrocytes. In neurons, extensive research highlights the importance of spatially organized cytoskeletal networks in establishing structural polarity and functional specialization across distinct cellular compartments. For instance, differences in microtubule polarity and post-translational modifications (PTMs) between axons and dendrites enable targeted transport of axonal versus dendritic cargo to support subcellular specification and function (Moutin et al., 2021; Iwanski and Kapitein, 2023; Kelliher et al., 2019). In addition, distinct actin microstructures and patterning further support axonal and dendritic specialization as well as synaptic function (Leterrier et al., 2017; Papandréou and Leterrier, 2018; Konietzny et al., 2017; Ouzounidis et al., 2023), while neurofilaments, neuron-specific IFs, maintain axonal caliber (Hoffman et al., 1987; Eyer and Peterson, 1994; Yuan et al., 2017). Thus, the importance of complex cytoskeletal regulation in neurons hints that the interplay between these three classes of cytoskeleton also underlies compartment-specific structure and function in astrocytes.

Unfortunately, fundamental questions about cytoskeletal organization in the branched processes of astrocytes have remained unresolved due to technical limitations. Resolving the fine detail of highly branched astrocyte processes *in vivo* via two-photon microscopy is technically challenging due to limitations in optical resolution and disentangling signal from noise in complex tissues. *In vitro* cell culture methods circumvent these issues and enable analysis of cell-autonomous mechanisms of cytoskeletal organization. However, traditional serum-based methods of culturing astrocytes produce flat, fibroblast-like cells with short protrusions and gene expression profiles resembling reactive astrocytes (Foo et al., 2011; Zhang et al., 2016). To address these limitations, we used immunopanning to isolate astrocytes from rodent brains. In immunopanning, a mixed population of cells is passed over a series of cell-surface antibodies to remove unwanted cell types and positively select a cell type of interest with high purity and cell viability. We cultured immunopanned astrocytes in serum-free, chemically defined conditions that promote complex branching and a stellate morphology, features reminiscent of healthy astrocytes *in vivo* (Zhang et al., 2016; Foo et al., 2011; Foo, 2013). To comprehensively visualize the astrocyte cytoskeleton in its native state, we leveraged immunostaining with confocal microscopy as well as cryo-ET to image cell structures with high resolution. Our work reveals distinct spatial organization of different elements of the cytoskeleton in the complex branches of astrocytes. The IF GFAP and microtubules define thicker main processes, while actin extends the boundaries of the cell through intricate microprocesses (e.g., filopodia, lamellipodia) at the cell periphery, and a novel structure that we call reticular webbing, which is found near the soma and connects thicker core branches together. Our observations uncover a spatially patterned cytoskeletal architecture that balances structural stability in proximal portions of branches with the capacity for functional specialization and dynamic remodeling at distal process tips.

## Results

### Immunopanned cultured astrocytes display complex branching morphology

We purified primary astrocytes from neonatal rat brain cortex using immunopanning, a method that selects for cells bearing astrocyte-specific cell surface markers and yields astrocyte cultures that are >98% pure and nearly devoid of neurons. Following published immunopanning protocols astrocytes were grown in serum-free media for two weeks (DIV14, days *in vitro*) to allow them to mature (Foo et al., 2011; Foo, 2013). The transcriptomes of immunopanned astrocytes resemble those of healthy astrocytes *in vivo* (Foo et al., 2011; Zhang et al., 2014, 2016). Similar to *in vivo* astrocytes (Fig. 1A), immunopanned astrocytes are highly ramified, with a complex stellate morphology visualized by GFAP and α-tubulin staining (Fig. 1B, S1A).

**Fig. 1.**
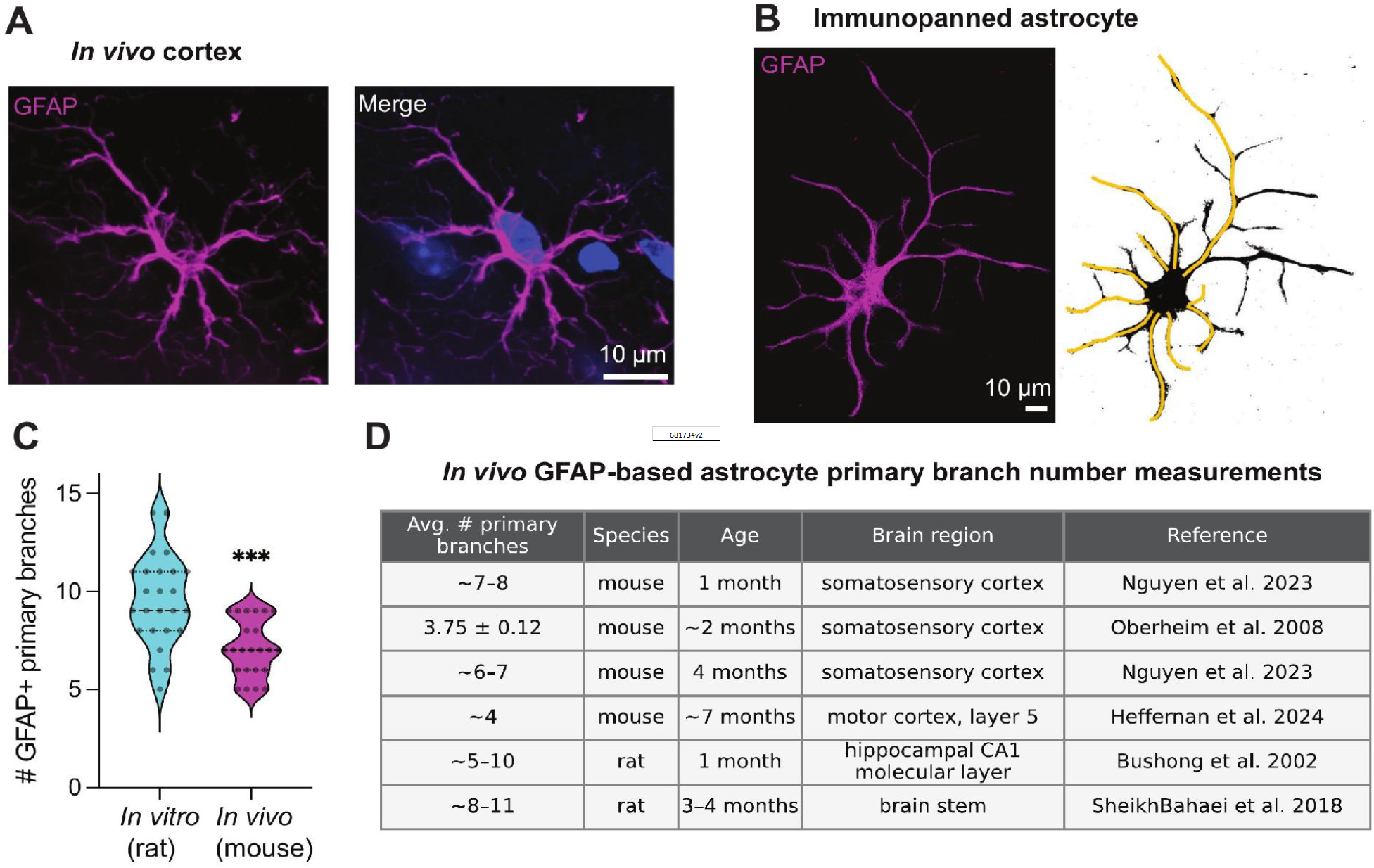
Immunopanned astrocytes exhibit GFAP-positive primary branch numbers comparable to astrocytes *in vivo*. **(A)** Representative image of a GFAP-positive astrocyte in mouse cortex. Sections were stained for GFAP to label astrocyte branches, DAPI to label nuclei, and CD31 to label blood vessels. An astrocyte endfoot in contact with a blood vessel is visible. Scale bar, 10 µm. **(B)** Representative immunopanned rat astrocyte (DIV14) cultured in serum-free conditions and stained for GFAP. Right, corresponding branch tracing illustrating GFAP-positive primary branches. Primary branches were defined as major GFAP-positive processes emerging from the edge of the soma. Scale bar, 10 µm. **(C)** Quantification of GFAP-positive primary branch number in immunopanned astrocytes compared with GFAP-labeled cortical astrocytes *in vivo*. Each dot represents one astrocyte; violin plots show the distribution of individual cells with median and interquartile range represented by dashed lines. Mean ± SEM for immunopanned cells was 9.5 ± 0.46. and for *in vivo* cells 7.1 ± 0.30. n = 25 immunopanned astrocytes from 3 independent cultures (from 3 litters of rats) and n = 23 *in vivo* cortical astrocytes from 3 mice. Mann-Whitney test, p < .0001. **(D)** Summary of published measurements of astrocyte primary/main branch number estimated from GFAP labeling in the rodent brain across other studies. These values are included to benchmark the proximal branch complexity of immunopanned astrocytes against prior *in vivo* measurements.

To compare *in vitro* to *in vivo* astrocyte morphology, we stained 4-month-old mouse cortex sections with GFAP (Fig. 1A). We traced GFAP-labeled branches from the soma edge to branch tip, classifying the longest continuous paths from the soma as primary branches. Immunopanned astrocytes average 9.5 ± 0.46 GFAP-positive primary processes per cell (Fig. 1C). This is comparable to our *in vivo* measurements in mouse cortex of 7.1 ± 0.30 GFAP-positive primary processes per cell (Fig. 1C). This modest difference is likely an expected consequence of the culture environment: isolated astrocytes lack the neighboring cells, neurons, vasculature, and three-dimensional matrix that impose contact-mediated tiling and constrain process extension *in vivo* (Stogsdill et al., 2017; Rodriguez Salazar et al., 2025; Gonzales et al., 2026; Tan et al., 2023). Thus, future co-culture experiments with astrocytes and neurons, for example, may more closely recapitulate *in vivo* measurements. Nevertheless, both measurements fall between the wide range (4–11 processes per cell) quantified by multiple *in vivo* rodent studies across different ages and brain regions (Heffernan et al., 2024; SheikhBahaei et al., 2018; Bushong et al., 2002; Nguyen et al., 2023; Oberheim et al., 2008) (Fig. 1D).

To characterize astrocyte branching architecture, we traced α-tubulin-labeled branches (Fig. 2A). We defined each secondary branch as the longest branch arising from a primary branch, and so forth (Fig. 2B). Tubulin labeling was used for branch architecture analysis because it resolves a fuller branching hierarchy, capturing more fine, short processes not consistently resolved by GFAP (Fig S1A–C). Immunopanned astrocytes displayed a mean of 10.0 ± 0.53 tubulin-positive primary branches per cell (Fig. 2C), with an average length of 48.0 ± 3.27 μm (Fig. 2D). Comparing different branch classes, we found that secondary branches are the most abundant (Fig. 2C) and that higher-order branches are progressively shorter (Fig. 2D). Together, these metrics reflect a hierarchical organization that mirrors the graded complexity of astrocyte arbors.

**Fig. 2.**
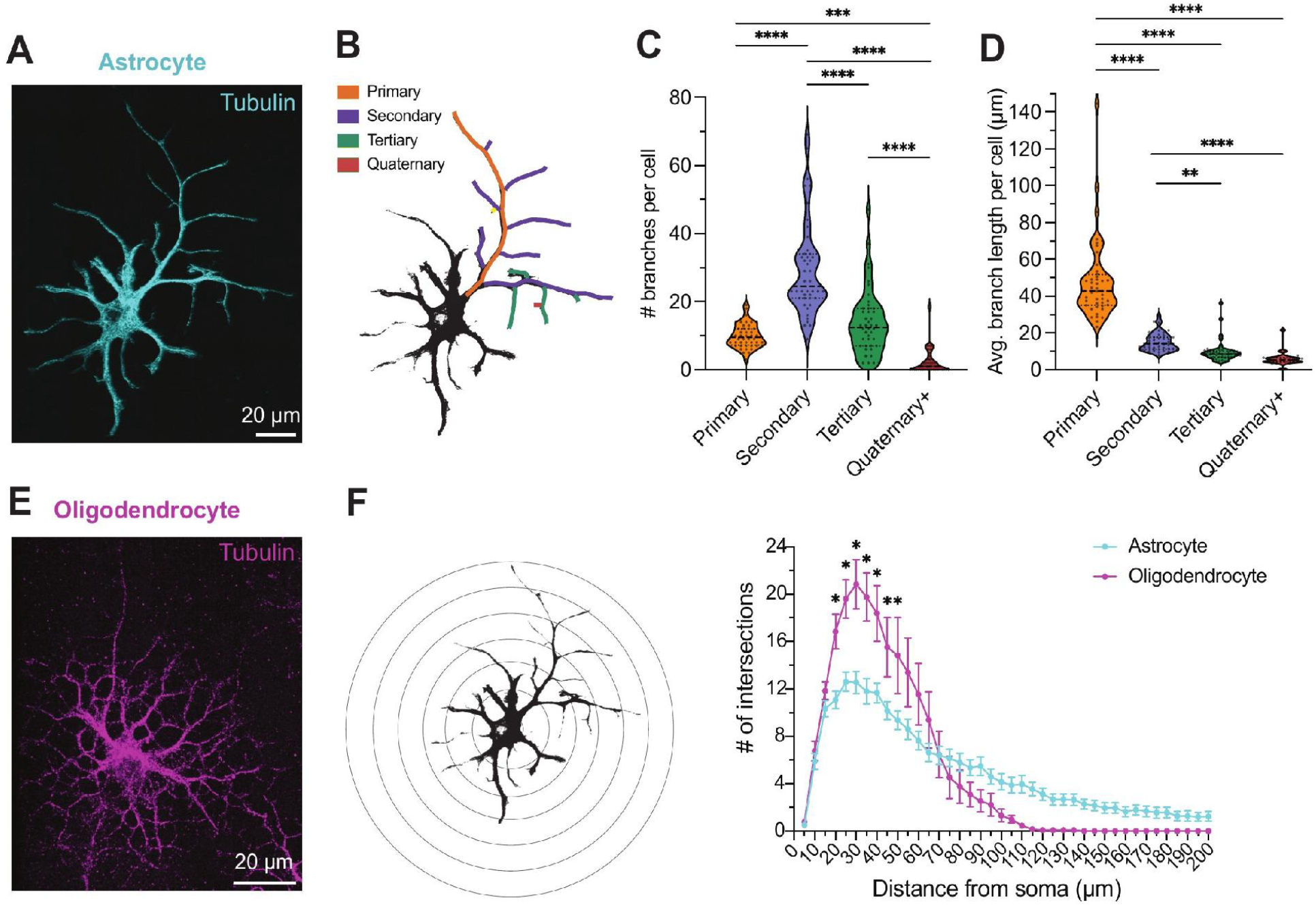
Immunopanned astrocytes exhibit a distinct, highly ramified microtubule architecture. **(A)** Representative immunofluorescence staining of tubulin (tyrosinated α-tubulin (YL1/2 antibody)) in immunopanned primary rat astrocyte at DIV14. **(B)** Trace of tubulin staining in (A) with color coding to illustrate branch order classification (primary, secondary, etc.) used throughout the study **(C)** Quantification of average number of branches per cell for each branch order based on tubulin staining in immunopanned rat astrocytes. Mean ± SEM for primary, secondary, tertiary, and quaternary branches, respectively, is 10.0 ± 0.52, 29.6 ± 2.1, 13.9 ± 1.45, and 2.5 ± 0.64. Friedman test with Dunn’s multiple comparisons. **(D)** Quantification of average branch length per cell for each branch order based on tubulin staining in immunopanned rat astrocytes. Mean ± SEM for primary, secondary, tertiary, and quaternary branches, respectively, is 48.0 ± 3.27, 15.1 ± 0.73, 9.8 ± 0.86, and 5.9 ± 0.75. Kruskal-Wallis test with Dunn’s multiple comparisons. In (C) and (D) violin plots show the distribution of individual cells with median and interquartile range represented by dashed lines. n = 44 cells from 4 independent cultures (from 4 separate rat litters), **(E)** Representative immunofluorescence staining of tubulin (tyrosinated α-tubulin (YL1/2 antibody)) in immunopanned DIV3 differentiated oligodendrocyte. **(F)** Quantification of Sholl analyses of astrocytes and oligodendrocytes, with example of an astrocyte on evenly spaced concentric rings for Sholl analysis pictured to the left of the graph. Astrocytes have significantly fewer branches than oligodendrocytes at distances between 20 µm and 50 µm away from the center of the soma. Mean ± SEM, n = 13–25 cells from 3–4 independent cultures per cell type (from 3–4 litters of rats). Mixed-effects model with repeated measures followed by Bonferroni multiple comparisons testing; * indicates p ≤ 0.05, ** p ≤ 0.01, *** p ≤ 0.001, **** p≤0.0001.

### Astrocyte branching morphology is unique from other glial cells

We also asked whether the microtubule branching pattern in astrocytes is distinct from that of other ramified glial cell types. Since microglia are much smaller (Nimmerjahn et al., 2005) and arise from a different lineage than other glial cells (Ginhoux et al., 2010), we compared astrocytes to immunopanned oligodendrocytes that were matured in differentiation media (Fig. 2E). Although the frequency and length of different branch orders (e.g., primary, secondary, etc.) was similar across cell types (Fig. S1D-E), we observed striking differences by Sholl analysis, which examines how branching complexity varies depending on distance from the soma or cell body. Oligodendrocytes have more proximal branches, peaking around 35 µm away from the cell body, and branch complexity drops off sharply after that, with few branches present >100 μm from the cell body. In contrast, astrocytes have significantly fewer proximal branches in the range of 20–50 µm away from the soma, their branching complexity declines more gradually, and branches are still present up to 200 μm away from the soma (Fig. 2F). Together, astrocytes and oligodendrocytes show broadly similar yet distinct tubulin branching patterns, suggesting they retain some intrinsic cell-type specificity in culture.

### Microtubule polarity is predominantly plus-end-out in the primary branches of immunopanned astrocytes

Microtubule polarity lays the groundwork for the directionality of motor-based cargo transport, which is particularly important in cells with long complex branches. To define microtubule polarity in morphologically complex immunopanned astrocytes, we expressed EB3-mNeonGreen in these purified cells to label the growing plus ends of microtubules (Fig. 3A) and performed live-cell imaging (Movie S1). We focused our kymograph analysis on EB3 runs in primary branches, where anterograde movement was defined as movement away from the soma. This analysis showed that microtubule polarity in immunopanned astrocytes was largely plus-end-out, with 89.9% anterograde runs per cell (Fig. 3B-C). Time-lapse images of EB3 movement highlighted EB3 localization at growing process tips (Fig. 3D-E) and potential nascent branch points (Fig. 3E, asterisks).

**Fig. 3.**
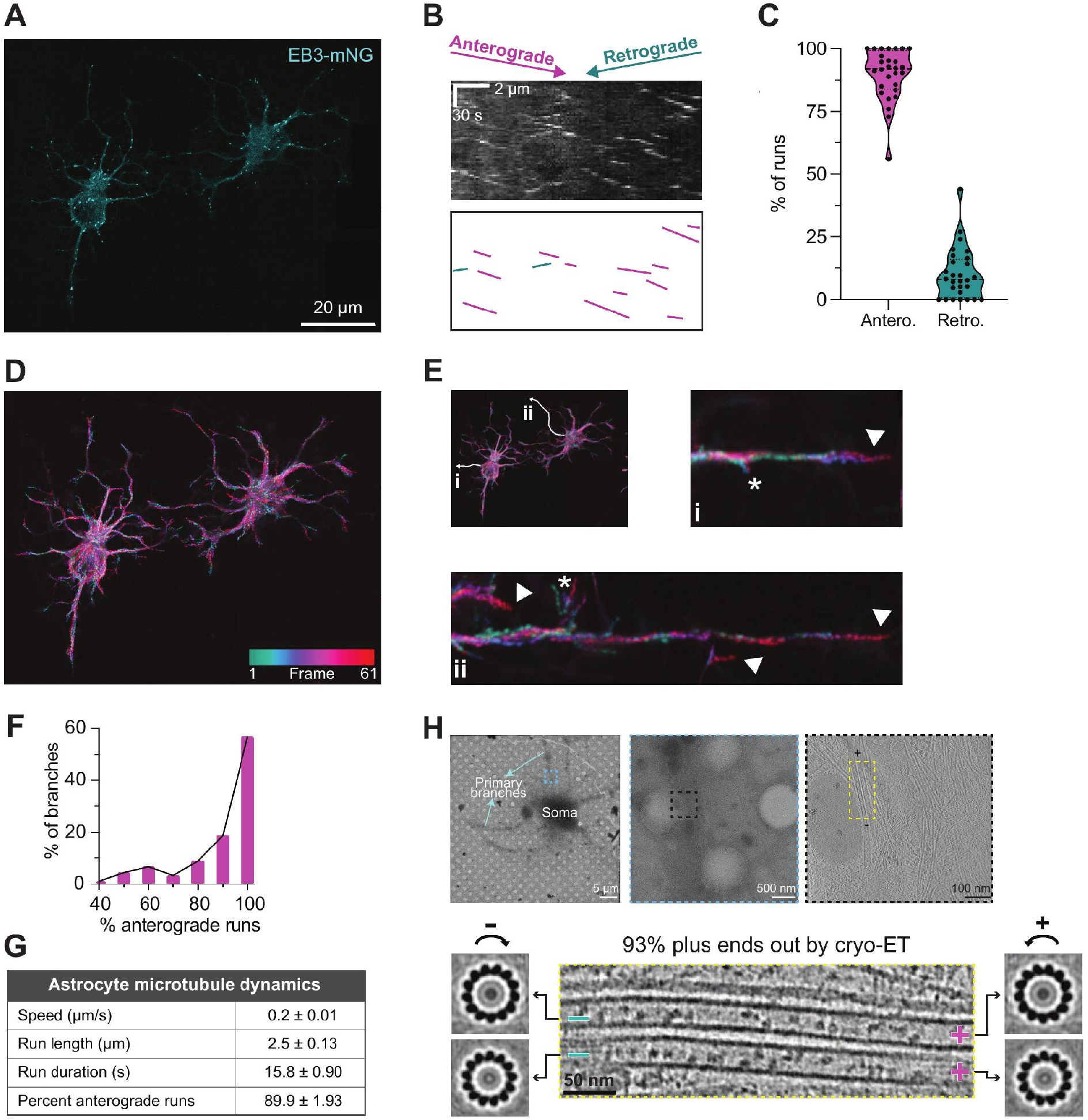
Microtubules in immunopanned astrocytes are predominantly oriented plus-end-out. **(A**) Single frame from a live-cell movie of DIV14 primary astrocytes expressing EB3-mNeonGreen (EB3-mNG) to mark the growing ends of microtubules. Processes from an adjacent astrocyte on the right were digitally erased for clarity. **(B)** Representative kymograph of one primary process from an EB3-mNG-expressing astrocyte, illustrating anterograde (purple) and retrograde (green) tracks. **(C)** Violin plot of the percentage of anterograde and retrograde EB3-mNG runs per cell. Violin plots are illustrated with lines representing median and dotted lines representing interquartile range (IQR). **(D)** Heat map of a time-lapse of a movie of EB3-mNG-expressing astrocytes in (A). EB3-mNG runs are represented as transitioning from green (beginning of movie) to red (end of movie). Frame rate was 3 seconds per frame; movie length was 3 minutes. **(E)** Magnified images of digitally straightened branches in (D), highlighting EB3-mNG localization at the ends of growing branches. Branches and magnified insets are labeled i and ii. Arrowheads label distal branch tips, with neighboring branch tips visible in ii. Asterisks label potential nascent branch points, defined as short microtubule branches with lengths of <5 µm. **(F)** Distribution of the percentage of anterograde and retrograde EB3-mNG runs per branch. **(G)** Table summarizing analysis of EB3-mNG runs in primary astrocytes, mean ± SEM (per cell). n = 110 kymographs from 28 cells. **(H)** Top left: example of a DIV13 astrocyte grown on a cryo-EM grid. The blue dashed box highlights the region of the primary process

We wondered whether all primary astrocyte branches were plus-end-out or if there were some astrocyte branches with mixed polarity, similar to polarity differences in axons versus dendrites (Yau et al., 2016). Thus, rather than calculating averages per cell, we binned events for each branch. We observed that most branches (87%) had predominantly plus-end-out polarity, with ≥70% anterograde runs.

However, a small fraction of branches (13%) had more mixed polarity with 40–70% anterograde runs (Fig. 3F). In addition to polarity, we determined the average speed, run length, and run duration per cell of EB3 comets (Fig. 3G, S2A–C).

We independently assessed microtubule polarity in immunopanned astrocytes using cryo-ET (Fig. 3H), which allowed us to view microtubules in their native state without the expression of an exogenous tagged protein. Analysis of cross sections from subtomogram averages revealed the polarity and protofilament number of microtubules in the primary processes. The radial skew (clockwise versus counterclockwise) of microtubule cross sections (Sosa and Chrétien, 1998) revealed that most microtubules (93%) were oriented with plus-end-out polarity (Fig. 3I). In addition, we observed most microtubules in primary processes contained 13 protofilaments (Fig. 3I and S2D), consistent with the canonical structure observed for neuronal microtubules (Foster et al., 2022; Atherton et al., 2018). The 13-protofilament geometry and plus-end-out polarity was maintained even in the presence of lattice breaks (Fig. S2D). Overall, these data indicate that microtubules in primary processes are highly ordered, both in terms of protofilament structure and polarity, consistent with a stably maintained microtubule architecture. used for acquiring a tomographic tilt series, Scale bar: 5 µm. Top middle: zoomed-in image of the primary process, highlighting the region of the tomogram using a black dashed box. Scale bar: 500 nm. Top right: 2D slice of a reconstructed tomogram highlighting the microtubules in the yellow dashed box used for determining polarity and protofilament number. Scale bar: 100 nm. **(I)** A zoomed in view of the two adjacent microtubules (center), and end-on view of their corresponding subtomogram averages showing 13-protofilament architecture and plus-end-out polarity, based on protofilament skew. Plus and minus ends are indicated. 93% of microtubules exhibited minus ends oriented toward the cell body and plus ends directed towards the tip of the process. n = 72 microtubules from 18 different primary branches and 9 different cells.

### Microtubules in astrocyte primary processes display markers of stabilization

One mechanism that facilitates microtubule stability is tubulin PTMs, which can modulate microtubule dynamics and functions such as cargo transport (Moutin et al., 2021; Janke and Magiera, 2020; McKenna et al., 2023). Thus, we analyzed tubulin detyrosination and acetylation (Fig. 4A), both of which have been associated with microtubule stabilization (Eshun-Wilson et al., 2019; Peris et al., 2009; Sanyal et al., 2023; Xu et al., 2017; Portran et al., 2017). Most primary branches in astrocytes were composed of microtubules exhibiting both acetylation and detyrosination (87%), with a significantly smaller percentage of primary branches displaying only acetylation (7%) or having neither modification (6%). In contrast, the proportion of secondary branches containing microtubules that were both acetylated and detyrosinated (43%) was only about half that of primary branches. The remainder of secondary branches contained microtubules that were either only acetylated (35%) or lacking either PTM (22%). Interestingly, we found no primary or secondary branches where detyrosination was present without acetylation (Fig. 4B). Thus, these results suggest that acetylation and detyrosination preferentially stabilize primary versus secondary branches in astrocytes.

**Fig. 4.**
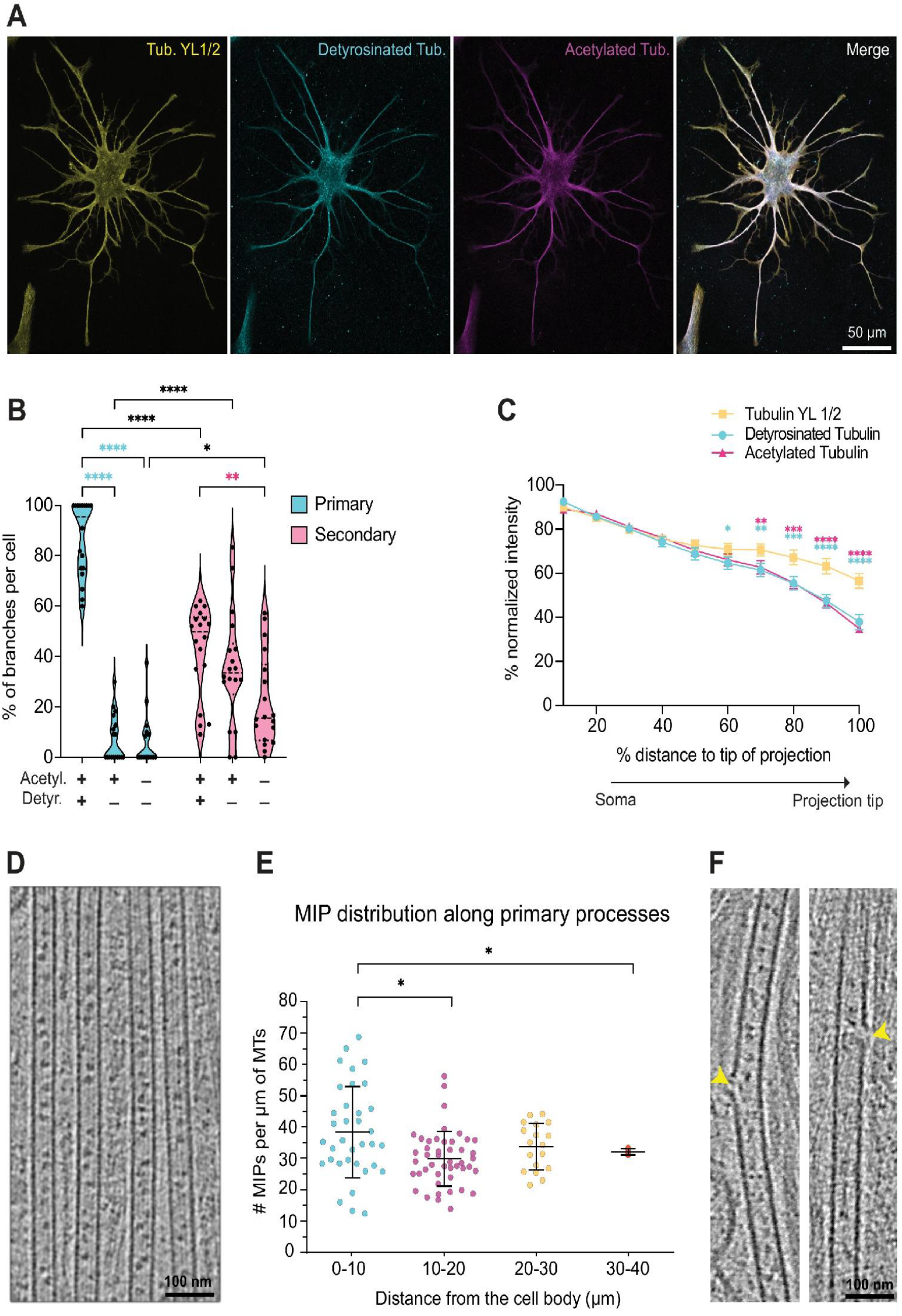
Microtubule-stabilizing PTMs and MIPs are present in the proximal regions of primary astrocyte branches. **(A)** Representative immunostaining for tyrosinated α-tubulin (YL1/2 antibody, yellow), detyrosinated tubulin (cyan) and both of which have been associated with microtubule stability (Cuveillier et al., 2020; Ichikawa et al., 2019; Rai et al., 2021), supports the idea that microtubules in the primary processes of astrocytes are highly stable. acetylated tubulin (magenta) in a DIV14 primary astrocyte. Merged image shows PTMs occur predominantly in primary branches with progressive loss of both PTMs toward distal tips. **(B)** Violin plots depicting percentages of primary and secondary branches positive for each PTM combination. Cyan asterisks represent comparisons within primary branches. Magenta asterisks represent comparisons within secondary branches. Black asterisks represent comparisons between primary and secondary branches. 2-way ANOVA with repeated measures followed by Tukey’s multiple-comparisons testing. Mean ± SEM of groups from left to right are 86.9 ± 3.6, 7.0 ± 2.2, 6.0 ± 2.4, 42.6 ± 4.2, 35.4 ± 5.3, and 22.1 ± 4.3%. **(C)** Fluorescence intensity of tyrosinated, detyrosinated, and acetylated tubulin along the length of primary branches. Both acetylated and detyrosinated microtubule intensities decrease significantly compared to tyrosinated microtubule intensities within the distal 30% of the branch. Fluorescence intensity was normalized against its maximum value per branch and distance is shown as a percentage of the total length of each branch (0% = edge of soma, 100% = projection tip). 2-way ANOVA with repeated measures followed by Dunnett’s multiple comparisons testing. Cyan asterisks show comparisons between tyrosinated tubulin and detyrosinated tubulin; magenta asterisks show comparisons between tyrosinated tubulin and acetylated tubulin. In (B) and (C), n = 16–18 cells from 3 independent cultures (from 3 litters of rats). * p ≤ 0.05, *** p ≤ 0.001, **** p ≤ 0.0001. **(D)** A 2D slice of a reconstructed tomogram showing the variable amounts of microtubule intraluminal proteins (MIPs) present in microtubules of DIV13 astrocyte primary processes. Scale bar: 100 nm. **(E)** Graph showing MIP frequency per µm of microtubule length in astrocytic primary processes, binned by distance from the cell body: 0–10 µm (n = 35 microtubules), 10–20 µm (n = 46), 20–30 µm (n = 18), and 30–40 µm (n = 4). Bars represent mean ± SD. One-way ANOVA with Tukey’s multiple comparisons. ** p < 0.01. **(F)** Representative examples of tomograms depicting microtubule lattice defects highlighted with yellow arrowheads. Scale bar: 100 nm.

We also examined whether levels of tubulin acetylation and detyrosination varied along the length of primary astrocyte branches. As astrocytes extend their processes, growing distal regions are likely newly formed and therefore more dynamic. Thus, we hypothesized that these PTMs are reduced at branch tips. In the proximal portion of primary branches (up to 50% of total branch length nearer to the soma), the relative fluorescence intensity of acetylated or detyrosinated tubulin was comparable to that of tyrosinated tubulin, which is detected by the α-tubulin (YL1/2) antibody (Wehland et al., 1984; Kilmartin et al., 1982). However, in the distal portion of branches (70–100% of total branch length), both acetylated and detyrosinated tubulin intensities were significantly lower than tyrosinated tubulin (Fig. 4C). Thus, the decline in the level of these PTMs in the distal primary processes may indicate that microtubules are less stable near branch tips.

Other structural features known to enhance microtubule stability are the presence of intralumenal proteins (MIPs) and lattice defects (Owa et al., 2019; van den Berg et al., 2023; Cuveillier et al., 2020; Ichikawa et al., 2019). Using cryo-ET, we observed that nearly all microtubules in astrocytic primary processes contain MIPs (Fig. 4D). The distribution of MIPs ranges from 15–70 per μm of microtubule with a modest enrichment of MIP frequency in the first 10 µm of primary processes (Fig. 4E). In addition to MIPs, we observed 1–4 defects per μm in the microtubule lattice (Fig. S2D-E); these defects range from the absence of a single tubulin dimer, to lattice breaks involving multiple dimers in one or more protofilaments (Fig. 4F). In contrast, lattice defects are much more rare in neurons, with one defect per 20 μm (Atherton et al., 2022). In all analyzed cases, the 13-protofilament geometry was maintained following a lattice defect (Fig. S2D), which has been interpreted to correspond to microtubule rescue events (Rai et al., 2021). Though protofilament transitions from 12 to 13 have been reported in individual microtubules in *Drosophila* neurites, they were not observed in mammalian neuronal axons (Foster et al., 2022). Thus, the high incidence of lattice defects in the absence of protofilament transitions suggests that astrocyte microtubules in primary branches display robust stability.

The resolution of the tomograms also allowed us to visualize the morphology of individual microtubule ends, where distinctive morphologies have been associated with growth and stability versus depolymerization (Kalutskii et al., 2025). We found that 18 out of the 34 microtubule ends analyzed displayed relatively short, flared morphologies, while the remaining 16 microtubule ends appeared blunt (Fig. S2F). Blunt and short flared ends have been observed for *in vitro* GMPCPP-bound microtubules and thus are thought to correspond to stable microtubules (Atherton et al., 2018). We did not observe ends with larger, curly protofilaments, which are associated with shrinking microtubules (McIntosh et al., 2018). The presence of MIPs and the prevalence of these microtubule end morphologies, both of which have been associated with microtubule stability (Cuveillier et al., 2020; Ichikawa et al., 2019; Rai et al., 2021), supports the idea that microtubules in the primary processes of astrocytes are highly stable.

### The cytoskeletal architecture of astrocytes is dominated by IFs

Interactions between different classes of cytoskeletal filaments play key roles in cellular physiology. To investigate this crosstalk in astrocytes, we examined the organization of cytoskeletal elements across scales, from whole-cell architecture to the nanoscale. Immunofluorescence revealed substantial overlap between the astrocyte-specific IF GFAP and tubulin in thicker primary branches. Actin was present throughout the cell and extended cellular boundaries beyond the microtubule-IF framework (Fig. 5A). Thus, the predominant method of using GFAP staining to visualize astrocytes misses a significant amount of the astrocyte cell space that contains actin.

**Fig. 5.**
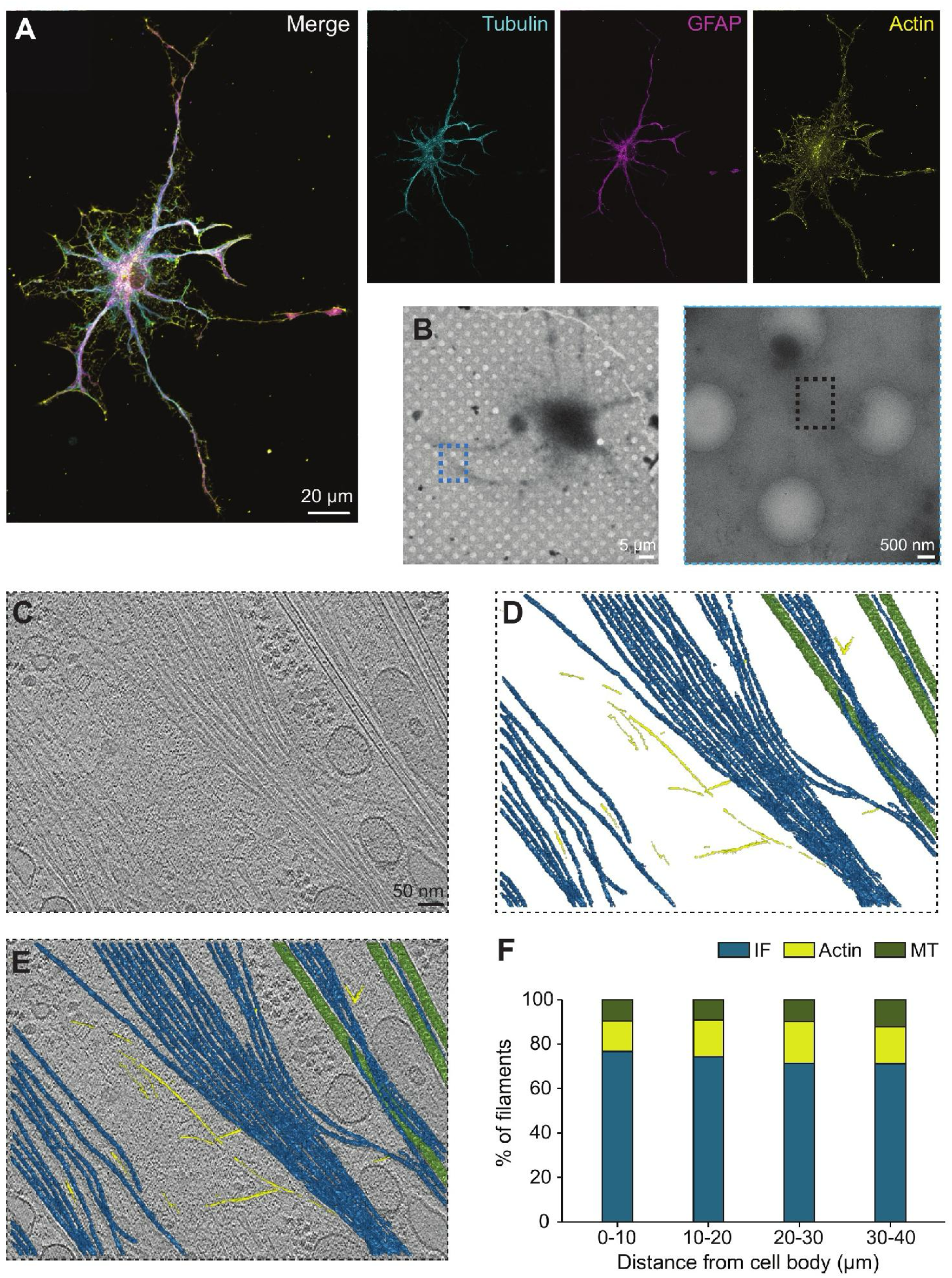
Abundant intermediate filaments define primary astrocyte processes while actin extends cellular boundaries. **(A-B**) Representative image of immunostaining for α-tubulin (YL1/2 antibody, cyan), the astrocyte-specific intermediate filament (IF) GFAP (magenta), and actin (phalloidin, yellow) in a DIV14 primary astrocyte. Processes from an adjacent astrocyte on the left were digitally erased for clarity. **(C)** 2D slice of a reconstructed and 3D CTF-corrected tomogram showing cytoskeletal filaments present in the primary process of a DIV13 astrocyte. Scale bar: 50 nm. **(D)** Trained neural network-based segmentation (on Dragonfly) of cytoskeletal filaments, with microtubules in green, IFs in blue, and F-actin in yellow. **(E)** Overlay of the segmented cytoskeletal features in (D) on the tomogram in (C). **(F)** Graph showing percent abundance of filament classes in astrocyte primary processes, divided into 10-µm bins from the edge of the cell body to 40 µm away. At the different locations analyzed, IFs comprise ∼70% of all filaments. The number of filaments represented in each bin are shown in Fig. S4A.

To visualize the interplay between these cytoskeletal systems at higher resolution, we applied cryo-ET (Fig. 5B) and used segmentation of our cryotomograms to highlight individual filaments (Fig. 5C-E). Using our tomograms, we estimated filament length in astrocyte primary processes, finding IFs and microtubules averaged ∼1.8 µm, whereas actin filaments were shorter, averaging ∼0.8 µm (Fig. S3A-B). Notably, we observed that astrocytes in the proximal 40 µm of primary processes contain a striking abundance of IFs compared to microtubules or actin, with IFs making up ∼70% of filaments (Fig. 5F, S3C). IFs outnumbered microtubules ∼7:1 and actin filaments ∼2:1 (Fig. S3D). The high ratio of IFs to microtubules in astrocyte primary processes resembles the distribution in medium to larger caliber myelinated axons, whereas microtubules dominate in dendrites (Hsieh et al., 1994; Hoffman et al., 1984; Reles and Friede, 1991).

The predominant IF expressed by astrocytes is GFAP, which forms both homopolymers and heteropolymers with another IF, vimentin (Chen and Liem, 1994; Eliasson et al., 1999; Potokar et al., 2020). Immunostaining of immunopanned astrocytes suggests strong overlap between GFAP and vimentin throughout the cell (Fig. S3E). However, because the ultrastructure of GFAP-vimentin heteropolymers has not yet been resolved, distinguishing GFAP from vimentin in tomograms was not possible. Therefore, we infer that the IFs seen by cryo-ET are likely a combination of GFAP homopolymers and GFAP-vimentin heteropolymers.

### The IF GFAP is enriched at astrocyte process tips

In neurons and other cell types, IFs interact with microtubules (Bocquet et al., 2009; Pradeau-Phélut and Etienne-Manneville, 2024; Schaedel et al., 2021). In our astrocyte tomograms, microtubules were frequently positioned closely alongside IFs in primary processes (Fig. 5E, S4A). To gain insight into the unexplored relationship between GFAP and microtubules in astrocytes, we examined their distribution in detail by immunofluorescence staining. Tubulin and GFAP signals largely colocalized along primary and secondary processes, but their intensities varied along the lengths of individual branches (Fig. 6, A–C). Notably, GFAP intensity was often more intense at process tips (e.g., Fig. 6B, Branches 8 and 9), though this was not universal across all branches (e.g., Fig. 6B, Branches 6 and 15). On average, GFAP intensity was significantly higher than tubulin intensity in the most distal 10 µm of processes (Fig. S4B).

**Fig. 6.**
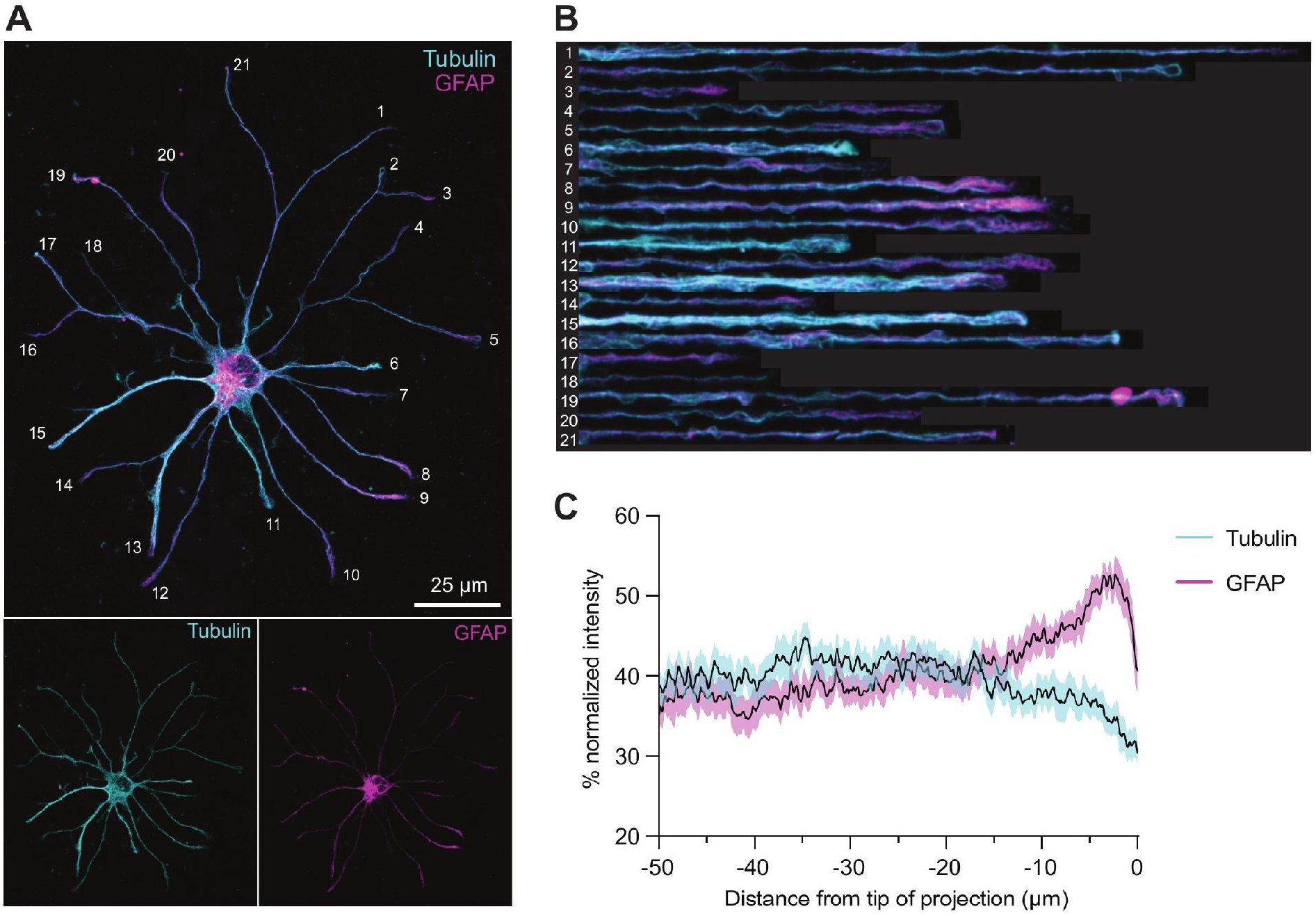
GFAP localization is enriched at distal process tips of primary astrocytes. **(A**) Representative image of immunostaining for α-tubulin (YL1/2 antibody, cyan) and GFAP (magenta) in a DIV14 primary astrocyte. **(B)** Images of digitally straightened processes labeled in (A), showing co-localization of tubulin and GFAP and distal enrichment of GFAP. **(C)** Fluorescence intensity of tubulin and GFAP across the distal 50 µm of branch tips shows an increase in GFAP intensity and slight decline in tubulin intensity within the most distal 15 µm of process tips of branches >50 µm. Fluorescence was normalized to its maximum value across the entire branch. n = 25 cells from 3 independent cultures. 5–13 processes were analyzed per cell.

Quantification across the distal 50 µm of processes revealed relatively stable levels of both filaments until ∼10–15 µm from the tip, where tubulin intensity declined and GFAP rose sharply (Fig. 6C). To better capture this shift, we calculated a distal enrichment ratio (average intensity across the most distal 10 µm divided by the average intensity over the preceding 40 µm). GFAP had a significantly greater distal enrichment ratio than tubulin (Fig. S4C). Across individual branch tips, 34% displayed strong GFAP enrichment, while 28% showed no enrichment. In contrast, only 6% of branch tips exhibited strong distal tubulin enrichment, while most tips (72%) showed no enrichment (Fig. S4D). Altogether, our results show that GFAP abundance frequently increases at the distal ends of long tubulin-positive processes, suggesting a potential specialized role for GFAP at process tips.

### Diverse actin microstructures in astrocytes: lamellipodia, filopodia, and reticular webbing

In addition to long, thick, shaft-like branches containing GFAP and microtubules, astrocytes have a fine meshwork of actin-rich microprocesses that significantly increases their surface area beyond the GFAP-positive skeleton (Fig. 5A and 7A). To visualize structures at the astrocyte periphery, we simultaneously stained for tubulin, GFAP, and actin, using phalloidin. These stains revealed a variety of actin microstructures and clear, consistent extension of actin localization beyond the main microtubule-IF framework. We classified actin structures as sheet-like lamellipodia, fine-tipped finger-like filopodia, or reticular webbing (Fig. 7, A-D), and also visualized them by cryo-ET (Fig. 7C).

**Fig. 7.**
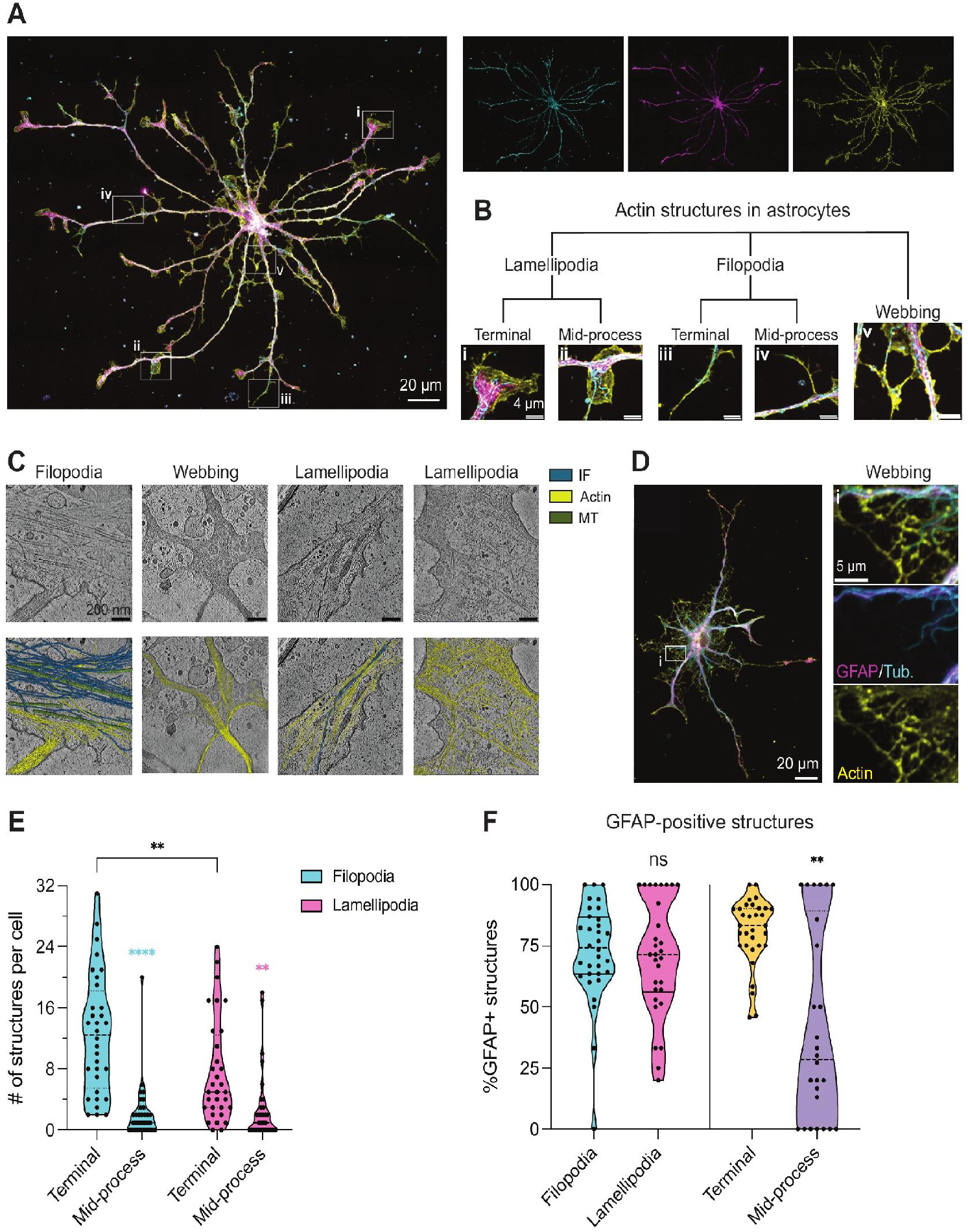
Diverse actin microstructures are present in the astrocyte periphery. **(A**) Representative image of immunostaining for α-tubulin (cyan), the intermediate filament GFAP (magenta), and actin (phalloidin, yellow) in a DIV14 primary rat astrocyte. Boxed areas are magnified in (B) and single channels are shown to the right. **(B)** Tree showing categorization of actin structures present in astrocytes, including terminal and mid-process filopodia and lamellipodia and webbing around the soma. Images are magnified from boxed areas of the cell in (A). All scale bars: 4 µm. **(C)** Top: example 2D tomogram slices of filopodia, lamellipodia, and webbing in DIV13 astrocytes. Bottom: overlay of segmentation of cytoskeletal filaments on the tomogram, with actin in yellow, microtubules in green, and IFs in blue. Scale bars: 200 nm. Additional example of actin webbing from the same cell pictured in Fig. 5A, with magnification of boxed region magnified to highlight distribution of the three cytoskeletal classes in the webbing. **(E)** Violin plots showing counts of filopodia and lamellipodia per cell, categorized by terminal versus mid-process location. On average, each astrocyte contained 12.6 ± 1.4 terminal filopodia, 2.2 ± 0.7 mid-process filopodia, 7.6 ± 1.2 terminal lamellipodia, and 2.8 ± 0.8 lamellipodia, mean ± SEM. 2-way ANOVA with repeated measures followed by Tukey’s multiple comparisons testing. **(F)** Violin plots showing the percentage of GFAP-positive filopodia versus lamellipodia (right) and terminal versus mid-process structures (left), Mann-Whitney tests. For (E and F), n = 33 cells from 4 independent cultures (from 4 litters of rats). ** p ≤ 0.01, *** p ≤ 0.001, **** p≤0.0001.

Reticular webbing is an intriguing structural feature in immunopanned astrocytes that. To our knowledge, has not been previously defined. Reticular webbing is an actin-based web-like network that is generally centered near the soma and links thicker tubulin- and GFAP-containing primary processes together (Fig. 7A-D, S5A). Most cells (85%) exhibited webbing, which radiated maximally outward, on average, 49.5 ± 5.4μm (mean ± SEM) away from the edge of the soma. Astrocytic reticular webbing consists of aligned actin filaments and often contains little detectable microtubule or IF signal by cryo-ET and immunofluorescence (Fig. 7C-D, S5A). In contrast, neuronal branchpoints visualized by cryo-ET have a mix of microtubules and unaligned actin (Nedozralova et al., 2022). Interestingly, small vesicles <100 nm in diameter are visible inside the reticular webbing (Fig. 7C).

We further quantified the relative distribution of filopodia and lamellipodia and their locations. On average, astrocytes had more filopodia than lamellipodia (Fig. S5B; 14.8 ± 1.78 filopodia vs 10.4 ± 1.83 lamellipodia per cell (mean ± SEM)). Lamellipodia and filopodia appeared both at branch termini and along branches as mid-process protrusions (Fig. 7, A-B), but were more abundant at termini. A majority of filopodia were terminal filopodia and a majority of lamellipodia were terminal lamellipodia (Fig. 7E, S5C-D). At mid-process sites, similar numbers and percentages of filopodia (8%) and lamellipodia (8%) were found (Fig. 7E, S5D). Overall, mid-process filopodia and lamellipodia are rarer than terminal ones.

Since IFs can interact with actin and influence its dynamics (Pradeau-Phélut and Etienne-Manneville, 2024), we hypothesized that GFAP may colocalize with actin structures. Indeed, the majority of filopodia (73%) and lamellipodia (71%) were GFAP-positive (Fig. 7F). We asked whether the presence of GFAP correlates with distance from the soma. Indeed, GFAP localization correlated with location along the branch; terminal structures were almost twice as likely as mid-process structures to be GFAP-positive (81% versus 42%). We also quantified filopodia and lamellipodia number per cell, segregating GFAP positivity and location (Fig. S5C). We found that, among terminal structures, GFAP-positive ones were located significantly farther from the soma than GFAP-negative ones (68 μm versus 42 μm) (Fig. S5E). Surprisingly, the magnitude of this difference (∼26 µm) was much greater than the average difference (∼10 µm) between terminal and mid-process structures (Fig. S5F) Overall, our data indicate that the majority of filopodia and lamellipodia, as well as most terminal actin structures are GFAP-positive.

### Disruption of actin polymerization alters primary branch architecture in developing astrocytes

Our data suggest that the three classes of cytoskeleton interact to build the distinct spatial patterning we observed in immunopanned astrocytes. Indeed, in neurons, the interplay between actin and microtubules is pivotal to developmental events, such as axon specification, axon elongation, and axon branching (Flynn et al., 2012; Bradke and Dotti, 1999; Dent and Kalil, 2001). To further interrogate the relationship between cytoskeletal classes in astrocyte branch morphogenesis, we asked how perturbing actin polymerization affects the architecture of primary branches, where IFs and microtubules provide a core structural framework. Thus, we treated DIV6 immunopanned astrocytes for 24 hours with 0.2 µM latrunculin A (LatA), a drug that sequesters actin monomers and promotes actin filament depolymerization. We chose DIV6, because it is an earlier developmental period with more active process protrusion and branching events than the more mature DIV13-14 timepoint used throughout the rest of our study. In LatA-treated cells, phalloidin staining confirmed marked reduction of filamentous actin, while GFAP and tubulin signals remained detectable and continued to mark primary processes (Fig. 8A). However, LatA-treated astrocytes displayed a significant increase in the number of tubulin-positive primary processes (Fig. 8B; LatA, 8.8 ± 0.30 vs. Veh., 6.6 ± 0.37 (mean ± SEM)) and average tubulin-positive primary branch length was significantly reduced (Fig. 8C-D; Veh., 43.1 ± 4.19 vs. LatA, 29.0 ± 2.19 µm (mean ± SEM)). Thus, perturbation of actin polymerization and dynamics led to more and shorter primary processes. Altogether, our data are consistent with a model in which actin, microtubules, and intermediate filaments make distinct contributions to the organization of complex astrocyte branches.

**Fig. 8.**
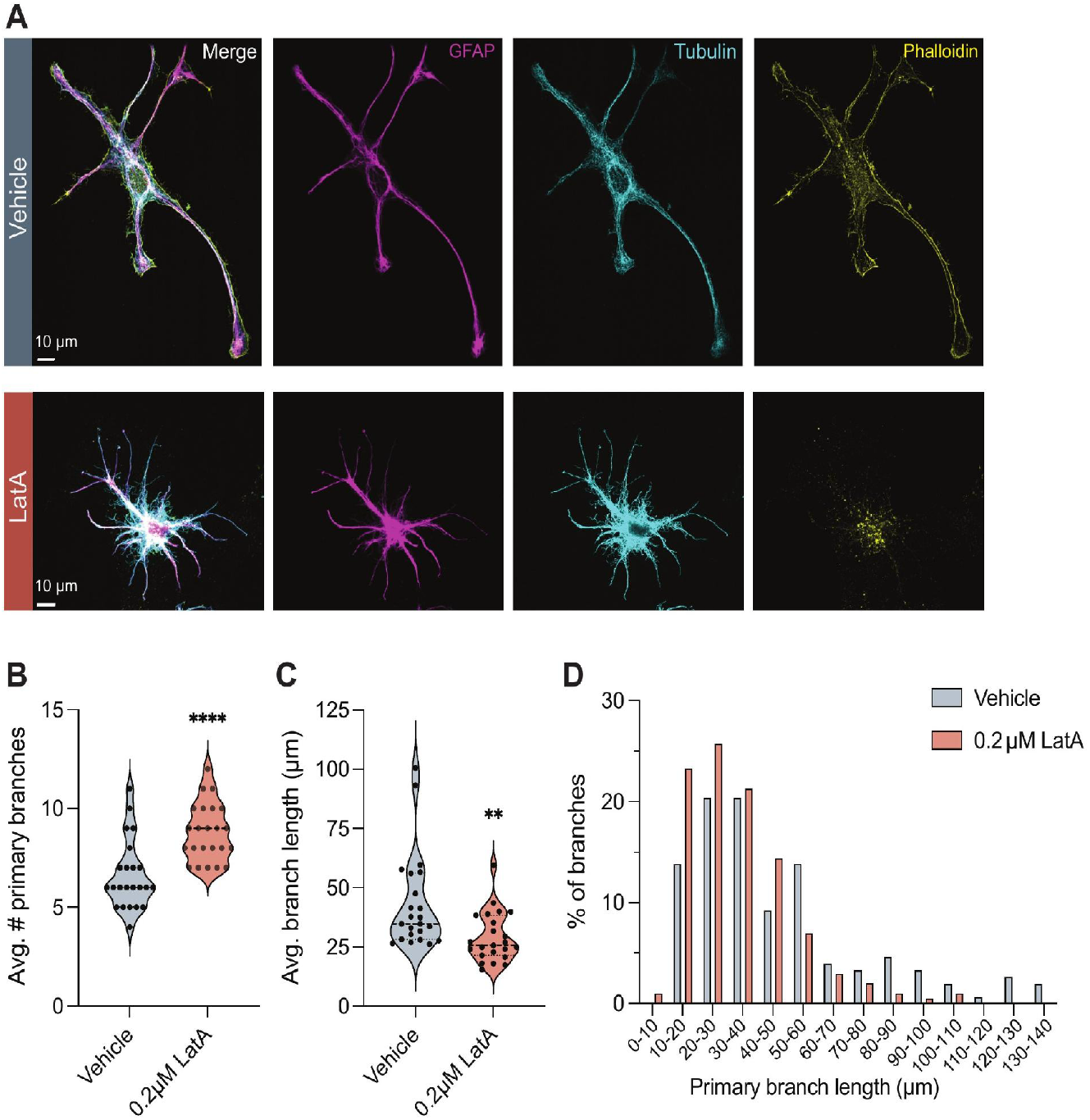
Acute actin disruption remodels primary branch architecture in developing astrocytes. **(A)** Representative immunofluorescence images GFAP, tubulin, and phalloidin staining in immunopanned astrocytes treated at DIV6 with vehicle or 0.2 µM Latrunculin A (LatA) for 24 hours and fixed at DIV7. Scale bars, 10 µm. **(B)** Quantification of average primary branch number per astrocyte, measured by tracing tubulin-positive branches, following vehicle or LatA treatment. Each dot represents one astrocyte; violin plots show the distribution of individual cells with median and interquartile range. **(C)** Quantification of average tubulin-positive primary branch length following vehicle or LatA treatment. LatA-treated astrocytes displayed reduced average branch length compared with vehicle-treated astrocytes. Each dot represents one astrocyte; violin plots show median and interquartile range. **(D)** Frequency distribution of primary branch lengths in vehicle-and LatA-treated astrocytes. LatA treatment shifted the distribution toward shorter primary branches. Data are from n = 23 vehicle-treated cells and n = 23 LatA-treated cells from 3 independent cultures. Frequency distribution represents 152 total branches from vehicle-treated cells and 202 total branches from LatA-treated cells (4–12 branches per cell). Statistical comparisons in (B) and (C) were performed using Mann–Whitney tests. **p ≤ 0.01, ****p ≤ 0.0001.

## Discussion

Our work provides compelling insights into the cytoskeletal architecture of complex astrocyte branches. Our results suggest a three-tiered arrangement of the cytoskeleton in astrocyte branches: (1) a stable inner core of acetylated and detyrosinated microtubules, further reinforced by IFs, (2) a transition zone where microtubules are less stable, and where enrichment of GFAP provides scaffolding to coordinate the microtubule and actin networks and support specialization of process tips, and (3) a morphologically diverse actin-based periphery that may be poised to respond to cues from neighboring cells and the environment. Altogether, our findings suggest that coordinated regulation of the cytoskeleton supports the structural and functional complexity of astrocyte branches.

### Implications for cargo transport in astrocytes

Our work informs future investigation of transport in ramified astrocytes and identifies two features of microtubule organization that are predicted to influence cargo transport. First, astrocyte microtubules in primary branches have a plus-end-out orientation, with 90% anterograde EB3 runs and 93% of microtubules being plus-end-out by cryo-ET (Fig. 3). This organization resembles the plus-end-out bias of axonal microtubules, although we have not directly measured cargo transport in this study. Our work establishes a robust system for future investigation into how plus-end-out-biased microtubule organization in core branches influences cargo movement in ramified astrocytes.

Second, astrocytic microtubule PTMs may also play a role in supporting cargo transport via microtubule stabilization and facilitation of the action of certain motors. We found that most primary processes (87%) contain detyrosinated and acetylated microtubules (Fig. 4A-B). In neuronal axons and cell lines, microtubule acetylation and detyrosination have been proposed to promote the selective binding to microtubules and motility of the anterograde motor kinesin-1 (Balabanian et al., 2017; Tas et al., 2017; Dunn et al., 2008; Reed et al., 2006; Cai et al., 2009). Indeed, primary astrocytes are known to transport a variety of cargos, including autophagosomes (Yuan et al., 2023; Kulkarni et al., 2020), lysosomes (Yuan et al., 2023), mitochondria (Jackson and Robinson, 2015), RNA granules (Meservey et al., 2021), and vesicles containing specialized astrocyte proteins, such as gap junction proteins and the glutamate transporter GLAST1 (Creighton et al., 2019). In these studies, many cargos moved in both anterograde and retrograde directions, but this work was largely performed in non-ramified astrocytes. We hypothesize the largely plus-end-out microtubule orientation we observed may support movement of diverse, specialized cargos toward distal branch tips, where astrocytes perform compartmentalized functions (e.g., water transport at interfaces with vasculature, or uptake of neurotransmitters near contact sites with synapses).

Intriguingly, the actin-rich reticular webbing centered around the soma that connects primary processes in astrocytes (Fig. 7, S5A) may also play a role in transport. By cryo-ET, this reticular webbing is often devoid of microtubules and IFs, but contains small vesicular organelles (Fig. 7C). Consistent with this, myosin-mediated transport has been observed in astrocytes (Stachelek et al., 2001) and membrane-bound organelles have also been found *in vivo* in very fine processes of astrocytes near synapses (Derouiche et al., 2015; Aboufares El Alaoui et al., 2020). Thus, these observations raise the intriguing possibility that myosin-mediated vesicular transport may occur along the actin filaments within the webbing.

### Stability of primary astrocyte processes

Astrocyte primary processes may utilize cytoskeletal mechanisms that depend primarily on microtubules and intermediate filaments to achieve structural stability. Our cryo-ET observations of primary astrocyte branches reveal abundant MIPs in the microtubule lumen (Fig. 4D-E), which are correlated with highly stable microtubules in neurons and microtubule doublets of cilia (Cuveillier et al., 2020; Owa et al., 2019; Ichikawa et al., 2019). Astrocyte microtubules also exhibit PTMs associated with microtubule stability, including acetylation (Eshun-Wilson et al., 2019; Xu et al., 2017; Portran et al., 2017) and detyrosination (Peris et al., 2009; Sanyal et al., 2023) (Fig. 4 A-C). Acetylation can alter inter-protofilament interactions and confer increased structural flexibility and resilience against mechanical stress (Eshun-Wilson et al., 2019; Peris et al., 2009; Sanyal et al., 2023; Xu et al., 2017; Portran et al., 2017). In cardiomyocytes, fibroblasts, and cell lines, detyrosination promotes binding of IFs to microtubules, which reduces depolymerization and increases the load-bearing capacity of detyrosinated microtubule bundles (Salomon et al., 2022; Kreitzer et al., 1999; Robison et al., 2016). Thus, the abundance of MIPs as well as of acetylated and detyrosinated microtubules may both contribute to branch stability in astrocytes, but further work is needed to dissect their precise functional roles.

Our surprising results showing no branches where detyrosination was present without acetylation (Fig. 4B) may suggest a temporal sequence of PTM acquisition in which acetylation precedes detyrosination during astrocyte maturation. Similarly, fibroblasts observed using super-resolution microscopy also did not harbor any non-acetylated-detyrosinated microtubules (Mohan et al., 2019). It is unlikely that these results in astrocytes and fibroblasts are due to antibodies competing for nearby binding sites, because acetylation occurs inside the microtubule lumen while detyrosination occurs on the microtubule surface. Though it is known that acetylation can occur rapidly on the timescale of minutes while detyrosination accumulates more slowly (Tang et al. 2023), it remains to be determined whether PTM kinetics and order of acquisition are related.

In addition to microtubules, astrocytes contain abundant IFs that likely also promote structural stability. *In vivo* support for a role for GFAP in structural stability comes from GFAP knockout mice, which are far less resilient to blunt-force mechanical trauma (Nawashiro et al., 1998). Similarly, in neurons, neurofilaments increase the viscoelastic properties of axons, providing a buffer against mechanical stress (Grevesse et al., 2015; Kornreich et al., 2016) In particular, larger caliber axons have greater resistance to compressive forces (Magdesian et al., 2012). Medium to larger caliber sensory and motor axons with diameters >3 µm have neurofilament:microtubule ratios ∼10:1 (Hsieh et al., 1994; Reles and Friede, 1991; Hoffman et al., 1984). In the primary processes of immunopanned astrocytes, we observe a high IF:microtubule ratio of ∼7:1 by cryo-ET (Fig. 5, S3D), which may indicate a similar role for IFs in maintaining process structural integrity in response to compressive and tensile stress. Synergy between the abundance of IFs and stabilizing microtubule PTMs may also enhance branch stability, as discussed above for detyrosination. Indeed, GFAP and tubulin colocalize along the lengths of most primary branches (Fig. 6) and an early study in non-ramified primary rat astrocytes observed co-localization of GFAP with acetylated microtubules (Cambray-Deakin et al., 1988).

Finally, the spatial organization of microtubule PTMs we observed may contribute to two important properties of *in vivo* astrocyte morphogenesis: dynamicity and tiling. Two-photon microscopy of astrocytes *in vivo* found that microtubules were mostly stable over 20-minute increments, but events of retraction, extension, and branching occurred at distal ends (Eom et al., 2011). These findings are consistent with our results showing that microtubule-stabilizing PTMs are not present at distal tips, with decreased abundance of acetylated and detyrosinated tubulin at the most distal third of primary processes (Fig. 4C). This spatial patterning is predicted to decrease microtubule stability at distal ends of processes, rendering them more permissive for local dynamic remodeling. Tiling is the process by which individual astrocytes establish and maintain distinct non-overlapping spatial domains within the brain (Bushong et al., 2002; Barber et al., 2021). As an astrocyte defines its territory from that of a neighboring astrocyte, branches may need to bend or twist to change directions. Acetylation imparts increased flexibility to microtubules, buffering against mechanical forces to allow bending (Xu et al., 2017; Portran et al., 2017; Janke and Montagnac, 2017). Indeed, in fibroblasts, microtubule acetylation has been linked to contact inhibition (Aguilar et al., 2014), a cellular mechanism that prevent cells from continually growing into one another. Thus, microtubule acetylation may also play an important role in astrocyte tiling by supporting mechanical flexibility of microtubules as branches survey the environment and find distinct territories to occupy.

### Interplay between classes of cytoskeleton in astrocyte morphology

While intermediate filaments and microtubules define the core scaffold of primary astrocyte branches, our LatA experiments demonstrate that actin also contributes to primary branch architecture. When we perturbed actin polymerization, we observed more microtubule primary processes. Disrupting actin polymerization also reduced average primary branch length by ∼33% (Fig. 8C-D), consistent with a role for actin in supporting elongated primary processes. This effect could reflect a contribution of actin to branch maintenance, continued branch extension, or both. These possibilities are not mutually exclusive and suggest that actin coordinates with microtubules and/or IFs to shape the boundaries and length of primary astrocyte branches.

Our result indicating that the interplay between different classes of cytoskeleton is important for establishing astrocyte branching patterns and morphology is consistent with classic observations in neurons. Similarly, actin depolymerization also affects neuronal process extension (Bradke and Dotti, 1999). Specifically, local severing of F-actin by ADF/cofilin is thought to promote the protrusion and bundling of microtubules, the backbone of new neuronal processes (Flynn et al., 2012). Actin dynamics are also important for axon branching and branch elongation. At neuronal growth cones, actin-based protrusion is thought to pioneer branch extension. Indeed, LatA treatment decreases the number of axon branches (e.g., secondary branches) and decreases axon branch length (Dent and Kalil, 2001) Though our work begins to dissect the contribution of actin polymerization in astrocyte branching, future work using pharmacological perturbation and genetic ablation will help dissect the specific contributions of individual actin regulators to branch formation and elaboration in astrocytes.

### Subcellular structures for specialized astrocytic functions

*In vivo*, astrocytes produce two types of processes that are morphologically and molecularly distinct, perisynaptic astrocyte processes (PAPs) and endfeet (Boulay et al., 2019, 2017; Soto et al., 2023; Salmon et al., 2023; Stokum et al., 2021). PAPs closely associate with neuronal synapses, buffer glutamate, and dynamically remodel in response to neuronal activity (Lawal et al., 2022; Henneberger et al., 2020). Endfeet are critical for maintaining the blood–brain barrier and coupling blood flow to neuronal activity (Heithoff et al., 2021; Mishra et al., 2024; Morales et al., 2022). The actin-rich structures that we observed in our cultures may be consistent with features of PAPs and endfeet, which we discuss in the following paragraphs.

The filopodia that we observed in our astrocyte monocultures may parallel precursor structures to PAPs, due to three shared properties. First, terminal filopodia are very abundant in cultured astrocytes (Fig. 7D), as are PAPs, with an *in vivo* estimate of thousands per rodent astrocyte (Bushong et al., 2002; Oberheim et al., 2009). Second, previous 3D reconstruction of astrocytes by FIB-SEM and SBF-SEM showed synaptic contacts preferentially occur at fine terminal branches (“leaflets”) and sites along astrocyte processes that constrict (Aboufares El Alaoui et al., 2020; Salmon et al., 2023; Aten et al., 2022). Similarly, filopodia in our cultured astrocytes are small, finger-like projections. Third, recent work demonstrates actin remodeling in astrocytes is crucial for structural plasticity of PAPs in response to neuronal activity (Henneberger et al., 2020). This parallels the role of axonal dendritic filopodia during neuronal development, which rapidly and repeatedly extend and retract in search of synaptic partners (Cohen-Cory, 2002). A similar need for the precise sensing mechanisms and dynamic remodeling of filopodia likely operates at PAPs. Together, these observations suggest that filopodia may be well suited to give rise to PAPs, based on their abundance, morphology, and an *in vivo* role for rapid actin remodeling.

In contrast to PAPs, endfeet are broad and sheet-like in order to fully cover the vasculature and therefore are more consistent with the lamellipodia and GFAP-rich terminals that we have identified. Notably, in human and rodent brains, GFAP is enriched in astrocyte endfeet (Oberheim et al., 2009). Indeed, GFAP is distally enriched in cultured astrocyte processes (Fig. 6) and a subpopulation of terminal actin structures colocalize with GFAP (Fig. 7). The large surface area of endfeet should enable more extensive coverage of the vasculature and a more stable adhesive surface to withstand repeated changes in vessel mechanical pressure. Because IFs can regulate focal adhesion dynamics and actomyosin contractility (Pradeau-Phélut and Etienne-Manneville, 2024; De Pascalis et al., 2018), distally enriched GFAP may tailor local actin responses to dynamic adhesive and mechanical forces, such as variable stiffness of the vasculature in response to vessel constriction and dilation. Thus, the interplay between GFAP and actin in distal astrocyte lamellipodia is especially poised to respond to vascular mechanical function. Future co-culture studies pairing astrocytes with neurons or vascular cells could assess whether actin-based filopodia and GFAP-rich lamellipodia contribute to the formation of PAPs and endfeet.

### Heterogeneity of astrocyte processes

A recent landmark study made the surprising observation that each astrocyte contacts at least one blood vessel. By two-photon microscopy, all of the ∼700 astrocytes analyzed in the mouse brain cortex contacted vasculature, with most astrocytes contacting 2–5 blood vessels (Hösli et al., 2022). This finding has important implications for astrocyte biology, as the proportion of astrocytes contacting a vessel was previously unknown and assumed to be only a smaller subset of cells. In other words, each astrocyte possesses heterogeneous branches – those that terminate in PAPs associated with neuronal synapses and those that terminate in endfeet associated with blood vessels. This is analogous to the compartmentalization of neuronal processes, where axons and dendrites adopt distinct morphologies and functions. One key difference between neuronal processes is microtubule polarity; while axons have nearly uniform plus-end-out polarity, dendrites have mixed polarity. In cultured neurons, axon specification is an intrinsically programmed process whereby a single neurite from an initial pool of sprouted plus-end-out neurites elongates into the axon, while other neurites gradually shift toward a more mixed polarity as they mature into dendrites (Baas et al., 1989). In astrocytes, we observed a small percentage of processes (13.3%) that display mixed microtubule polarity (40–70% plus-end-out) (Fig. 3F). It is intriguing to speculate whether this subpopulation of mixed-polarity astrocyte processes might be functionally distinct, poised to give rise to either PAPs or endfeet. Alternatively, signals from neurons and the vasculature may be required for full maturation of PAPs and endfeet. Future experiments comparing monoculture, neuron coculture, vascular coculture, and astrocytes *in vivo* will be needed to determine precisely how intrinsic and extrinsic cues influence the specification of heterogeneous processes in astrocytes.

In conclusion, our findings lay the groundwork for further investigation into how cytoskeletal architecture and dynamics underpin astrocyte morphology and function. As a cellular system to investigate mechanisms of cytoskeletal organization, astrocytes present many fascinating and novel cell biology questions, including the complex interactions between multiple classes of cytoskeleton, the actin-rich reticular webbing that connects primary processes, and the ontological origin of process heterogeneity. Many concepts in neuronal cell biology, including cargo transport and local translation, likely play unexplored roles in establishing subcellular specificity and function in astrocyte processes (Meservey et al., 2021; Mazaré et al., 2021; Kemal et al., 2022). Moreover, these questions in astrocyte cell biology will have important implications in our understanding of the roles astrocytes play in neurological disease and open the door to exciting new topics of investigation

## Acknowledgements

We thank Drs. Antonina Roll-Mecak, James Hurley, and Matthew Welch for insightful discussions and Dr. Patricia Grob for technical assistance. We thank the Bay Area Cytoskeleton Symposium, which made this collaboration possible.

## Author contributions

Conceptualization: M.E.W., W.E.B., D.G., E.N., M.M.F. Analysis and visualization: M.E.W., W.E.B., D.G., N.J., J.J.B., E.A.L. Funding acquisition: E.N., M.M.F. Investigation: M.E.W., W.E.B., D.G., L.T.H., E.A.L. N.J. Supervision: M.E.W., W.E.B., E.N., M.M.F. Writing - original draft: M.E.W., W.E.B, D.G., M.M.F. Writing - review & editing: M.E.W., W.E.B., D.G., N.J., J.J.B., L.T.H., E.A.L., E.N., M.M.F.

## Materials and Methods

### Animals

Wild type C57BL/6J mice used for immunostaining of cortical astrocytes *in vivo* were obtained from The Jackson Laboratory (Strain #: 000664) (RRID:IMSR_JAX:000664). Experiments were performed using 4-month-old mice of both sexes. All primary cell experiments were conducted using Sprague Dawley CD/SD rats (Charles River Laboratories, Strain Code: 001) (RRID:RGD_734476). Litters were sacrificed between P5–P7 (postnatal day) by rapid decapitation. Cortices used for cultures were combined by litter, generating cultures of mixed sex. Rats and mice were housed on a 12-hour light/dark cycle with food and water *ad libitum*. All animal procedures were performed in accordance with protocols approved by either the University of California, Berkeley (AUP-2023-06-16512) or the National Institute of Neurological Disorders and Stroke Animal Care and Use Committee.

### Astrocyte immunopanning and culture

Methods for isolating and culturing immunopanned neonatal rat astrocytes were performed according to previous protocols (Foo, 2013; Foo et al., 2011) with minor modifications.

The day before immunopanning, 15 cm Petri dishes were coated by adding 60 µL secondary antibody, or 100 µg BSL-1 lectin (Vector Laboratories, L-1100-5), to 20 mL of 50 mM Tris-HCl (pH 9.5), ensuring the entire dish surface was covered, and incubated overnight at 4 °C. Secondary antibodies used were goat anti-mouse IgG + IgM, goat anti-mouse IgM μ-chain specific, and goat anti-rat IgG (Jackson ImmunoResearch, 115-005-044, 115-005-020, 112-005-167). The following day, the secondary-antibody solution was discarded and plates were washed 3 times with PBS. 20 μL of each primary antibody was added to 12 mL of a DPBS solution with 0.2% BSA (Sigma, A4161) and DNase (Worthington, LS002007); this solution was incubated on the same plates at room temperature until needed for immunopanning steps.

Following preparation of antibody-coated plates, cortices from 8 P5–P7 rats with meninges removed were minced into ∼1 mm^3^ pieces, split evenly between two 6-cm petri dishes, and digested for 45 minutes at 34 °C in 20 mL of enzymatic solution containing 0.0040 g _L_-cysteine (Sigma, C7477), 100 U (unit) papain (Worthington, LS003126), and 100 μL of 0.4% DNase. The papain solution was pre-equilibrated with carbogen (5% CO_2_, 95% O_2_) and carbogen was gently bubbled over cortices during digestion. The tissue was then washed 3X and triturated in low-ovomucoid solution to create a single cell suspension. Low-ovomucoid solution consisted of a pre-equilibrated buffer solution (0.46% _D_(+)-glucose and 26 mM NaHCO_3_ in Earl’s Balanced Salt Solution) containing 8.5 µM BSA (Sigma, A8806), 20 µM ovomucoid-based trypsin inhibitor (Worthington, LK003182), and 0.45 µM insulin (Sigma, I6634). Pre-equilibrated high-ovomucoid solution (21.2 µM BSA, 50 µM trypsin inhibitor) was carefully layered under the cell suspension in low-ovomucoid solution and the cells were spun down for 5 minutes at 120*g* at room temperature to fully quench papain activity. The cell pellet was then resuspended in panning buffer (0.02% BSA/DNAse and 0.45 µM insulin in DPBS) and filtered through a 40 µm cell strainer to remove clumps of cells and to generate a single-cell suspension. From this mixture, macrophages, microglia, endothelial cells, and oligodendrocyte precursor cells were removed by immunopanning with anti-mouse IgG+IgM secondary antibody, BSL-1 lectin, anti-CD45 antibody (BD Biosciences Pharmingen, 550539), and O4 hybridoma supernatant (homemade), respectively. Lastly, astrocytes were positively selected for using anti-ITGB5 antibody (eBioscience, 14-0497-82). Resulting astrocytes were lifted with 400 U trypsin (Sigma, T9935) in EBSS pre-equilibrated at 37 °C with 10% CO_2_ in the cell culture incubator. Astrocytes were then spun down in 30% heat-inactivated FBS (Gibco, 16000044) for 5 minutes at 130*g* at room temperature to quench trypsin activity. This was the only time during which cells were exposed to serum.

Astrocytes were then resuspended in growth medium and plated onto 12-mm sonicated glass coverslips pre-treated with 10 µg/mL poly-D-lysine (PDL) hydobromide (Sigma, P6407) in a 24-well cell culture plate, at a density of 20,000–40,000 cells/well. Cells were grown at 37 °C in 10% CO_2_ for 13–14 days in astrocyte growth medium containing 1:1 DMEM:Neurobasal mixture (Gibco, 11960-051, 21103-049) containing 100 U/mL penicillin/streptomycin (Gibco, 15140-122), 1 mM sodium pyruvate (Gibco, 11360070), 1X GlutaMAX (Gibco, 35050061), 1X DMEM-based SATO supplement (100 µg/mL BSA (Sigma, A4161), 100 µg/mL transferrin (Sigma, T1147), 16 µg/mL putrescine dihydrochloride (Sigma, P5780), 60 ng/mL progesterone (Sigma, P8783), and 40 ng/mL sodium selenite (Sigma, S5261)), 5 ug/mL N-acetyl-L-cysteine (Sigma, A9165), and 5 ng/mL HbEGF (Sigma, E4643). This media was refreshed every 4–7 days.

### Oligodendrocyte immunopanning and culture

Oligodendrocyte precursor cells (OPCs) were purified from neonatal Sprague-Dawley rat pups (P6–P8) (Charles River) of both genders by immunopanning (Dugas and Emery, 2013). Briefly, cortical tissue was dissociated by papain digestion and filtered through a 40-µm cell strainer to obtain a single-cell suspension. This suspension was incubated in two negative-selection plates, one coated with anti-RAN-2 hybridoma supernatant and the other coated with anti-GC (glucocerebrosidase) hybridoma supernatant, then in a positive-selection plate coated with anti-O4 hybridoma supernatant. All hybridoma supernatants were made in-house. Adherent cells were trypsinized and plated on 12-mm glass coverslips that were cleaned via sonication for 1 hour in ethanol, then coated with 10 µg/mL poly-D-lysine (PDL) hydrobromide at a density of 10,000–20,000 cells each. Cells were cultured at 37 °C and 10% CO_2_ in proliferation media containing PDGF (STEMCELL Technologies, 78095.1) and NT3 (STEMCELL Technologies, 78074), or differentiation media containing NT3, CNTF (STEMCELL Technologies, 78010.1), and T3 (Sigma, T6397). Immunopanned oligodendrocytes are >99% pure and express markers of myelinating oligodendrocytes following differentiation (Dugas et al., 2006; Zhang et al., 2014; Dugas and Emery, 2013).

### Immunofluorescence staining of mouse brain tissue sections

Mice were anesthetized with a ketamine/xylazine cocktail and euthanized by rapid cervical dislocation, followed by transcardial perfusion with ice-cold PBS. Brains were collected and post-fixed in 4% paraformaldehyde overnight at 4°C, then cryoprotected in 30% sucrose until embedding in Tissue-Plus O.C.T. Compound for sectioning. Coronal sections (40 μm) were cut on a Leica cryostat and stored free-floating in glycerol-based cryoprotectant at −20°C until staining. Sections containing both hippocampus and thalamus were selected for imaging and analysis of cortical astrocytes.

For staining, sections were washed 3 × 10 min in PBS-T at room temperature on a rocker, then incubated overnight at 4°C with primary antibodies diluted in staining buffer (1% donkey serum in PBS-T). The following day, sections were washed 3 × 10 min in staining buffer at room temperature, incubated for 2 h at room temperature with secondary antibodies, and mounted with ProLong Gold Antifade Mountant (Invitrogen). Z-stacks (0.2 μm step size) were acquired on a spinning-disk confocal microscope. Slides were imaged within 3 days from mounting using 60X or 100X objectives on a spinning-disk confocal (Nikon Eclipse-T2 inverted microscope with Yokogawa X1 spinning disk) with a sCMOS camera (Hamamatsu Orca-FusionBT).

### Immunofluorescence staining of cultured primary glial cells

All stainings besides experiments with LatA were performed on astrocytes fixed at DIV13-DIV14. LatA-treated astrocytes were fixed on DIV7 following 24 hours of 0.2µM LatA (Sigma, 428021) or DMSO vehicle treatment. Cultures were washed with 37 °C PHEM (60 mM PIPES, 25 mM HEPES, 10 mM EGTA, 2 mM MgSO_4_, pH 6.9–7) or PBS, and then fixed at 37 °C for 10 minutes with either fresh 4% PFA (Electron Microscopy Sciences, 15711) in PHEM containing 4% sucrose, or fresh 4% PFA in PBS. Then, coverslips were washed twice with PHEM or PBS, and permeabilized with 0.1% Triton X-100 in PHEM or PBS for 3 minutes. Following permeabilization, coverslips were washed 3X with PHEM or PBS, and then incubated in PBS-based (5% donkey serum (Sigma, D9663), 1% BSA (Sigma, A8806) in PBS) or PHEM-based blocking buffer (5% donkey serum, 1% BSA, 0.3 M glycine in PHEM) for 60 minutes at room temperature. Coverslips were incubated with primary antibodies diluted in blocking buffer overnight at 4 °C. The following day, coverslips were washed 3X with PHEM or PBS, then incubated with secondary antibodies and/or Phalloidin-647 diluted in blocking buffer for 1 hour at room temperature with protection from light. Then, coverslips were washed 3X with PHEM or PBS and mounted to slides using non-hardening Vectashield Plus (Vector Laboratories, H-1900). The use of PHEM vs. PBS buffers did not affect the fixation or staining quality of cytoskeleton and biological replicates were always exposed to the same buffer within individual experiments.

Slides were imaged on the same day or the following day using 60X or 100X objectives on a spinning-disk confocal (Nikon Eclipse-T2 inverted microscope with Yokogawa X1 spinning disk) with a sCMOS camera (Hamamatsu Orca-FusionBT). Z-stacks (0.5 μm step size) were acquired on a spinning-disk confocal microscope.

Primary antibodies and dyes (with dilutions):

Tyrosinated a-Tubulin YL1/2 (ThermoFisher, MA1-80017, 1:500) (RRID:AB_2210201)

Detyrosinated Tubulin (Abcam, ab48389, 1:200) (RRID:AB_869990)

Acetylated Tubulin (Sigma, T7451, 1:200) (RRID:AB_609894)

GFAP (Cell Signaling, 12389, 1:400) RRID:AB_2631098)

GFAP (Invitrogen, 13-0300, 1:200) (RRID:AB_86543)

Phalloidin-647 (Sigma, A30107, 1:400)

Vimentin (Abcam, ab24525, 1:400) (RRID:AB_778824) Secondary antibodies (with dilutions):

DyLight™ 405 AffiniPure® Donkey Anti-Rat IgG (H+L) (Jackson ImmunoResearch, 712-475-153, 1:250) (RRID:AB_2340681)

Alexa Fluor® 488 AffiniPure® Donkey Anti-Mouse IgG (H+L) (Jackson ImmunoResearch, 715-545-151, 1:500) (RRID:AB_2341099)

Alexa Fluor® 488 AffiniPure® Donkey Anti-Rabbit IgG (H+L) (Jackson ImmunoResearch, 711-545-152, 1:500) (RRID:AB_2313584)

Rhodamine Red™-X (RRX) AffiniPure® Donkey Anti-Mouse IgG (H+L) (Jackson ImmunoResearch, 715-295-151, 1:500) (RRID:AB_2340832)

Rhodamine Red™-X (RRX) AffiniPure® Donkey Anti-Rabbit IgG (H+L) (Jackson ImmunoResearch, 711-295-152, 1:500)

Rhodamine Red™-X (RRX) AffiniPure® Donkey Anti-Rat IgG (H+L) (Jackson ImmunoResearch, 712-295-153, 1:500) (RRID:AB_2340676)

Alexa Fluor® 647 AffiniPure® Donkey Anti-Rabbit IgG (H+L) (Jackson ImmunoResearch, 711-605-152, 1:500) (RRID:AB_2492288)

### Live-cell imaging

Immunopanned astrocytes used for live imaging were cultured on 35-mm glass-bottom dishes (Mattek, P35G-1.5-10-C) coated with PDL hydrobromide. Cells were seeded at a density of ∼60,000 cells per dish, transfected with EB3-mNeonGreen on DIV9 - 10, and imaged on DIV13 - 14. Transfection was performed with Lipofectamine 3000 (Invitrogen, L3000-001) according to manufacturer instructions, using 0.5 µg of plasmid per dish. EB3-mNeonGreen was a gift from Dorus Gadella (Addgene plasmid #98881; http://n2t.net/addgene:98881; RRID:Addgene_98881).

For live-cell imaging, the DMEM:Neurobasal base of astrocyte growth media was replaced with FluoroBrite DMEM (Gibco, A1896701) to limit background fluorescence. The cells were imaged in an environmental chamber at 37 °C with 5% CO2, and imaging was limited to one hour or less per plate to limit phototoxicity. Images were collected every 3 s for 3 min on a spinning-disk confocal (Nikon Eclipse-T2 inverted microscope with Yokogawa X1 spinning disk) with a sCMOS camera (Hamamatsu Orca-FusionBT). Movies were acquired within a single plane.

### Immunofluorescent image analysis

All image quantification was performed using the Fiji image analysis software (Ver. 2.16.0; Build 26d66057dd). Fluorescent intensities of tubulin PTMs, GFAP, and phalloidin were quantified by tracing each process with the Segmented Line Selection tool (20-pixel width), then exporting the data from the Plot Profile tool. The relative intensities of different stains were compared against one another by expressing fluorescent intensity values along individual processes as a percentage of the maximum intensity value of each marker along the process. These values are denoted as “% normalized intensity” in figures. Linearized representations of processes were generated by using the Straighten tool. For quantification and representation of tubulin PTMs as a percentage of total process length, mean values were binned into deciles.

### Sholl analysis

Images of tubulin-stained oligodendrocytes and astrocytes were traced using the Adobe Photoshop brush tool with a 5-pixel round tip for conversion into binary images. The freehand tool in Fiji was used to trace the cell body and to calculate a center of mass, which was used subsequently as the coordinates for the center of the cell. These coordinates and binary images were then input into the Sholl Analysis plugin (SNT 4.2.1) in Fiji and analyzed with a radius step size of 5 μm.

### Quantification of process number and length

The lengths of astrocyte branches were calculated by tracing tubulin- or GFAP-immunostained processes with the segmented line tool in Fiji. Primary processes were defined as a process extending directly from the edge of the cell body. Length was defined as the longest continuous path along the primary process and measured from the distance between the cell body edge and the process tip. Processes branching off from a primary process were defined as secondary processes. Tertiary processes were defined as those branching off from secondary processes, while quaternary processes were defined as those extending off of tertiary processes. Secondary, tertiary, and quaternary processes were traced from the base of the initial branch point out to the distal process tip. In cases where multiple continuous paths could be traced, the longest continuous path was assigned to the lowest order branch.

### Kymograph generation and microtubule polarity analysis

Kymographs were generated from live-cell images of EB3-mNeonGreen-transfected astrocytes using the Multi Kymograph tool in Fiji. Up to four kymographs from each cell were analyzed. The length and angle of each EB3 comet was measured using the Segmented Line tool and used to calculate direction, velocity and net displacement of each run. Two to four kymographs from each cell were analyzed. Kymographs were taken from predominantly primary processes with a smaller percentage (∼15%) taken from secondary processes. Average velocity, displacement, and percentage of distally- and proximally-traveling runs were calculated for each cell. Temporal heat maps of EB3 displacement were generated using the Temporal-Color Code tool in Fiji.

### Actin microstructure classification and quantification

Actin microstructures on tubulin-, actin-, and GFAP-immunostained astrocyte processes were classified and counted in Fiji based on the presence (filopodia) or absence (lamellipodia) of thin, pointed actin shapes protruding from the tip of the microstructure. We also quantified the location of filopodia and lamellipodia along processes (terminal or mid-process) and their distance from the cell body. Filopodia or lamellipodia microstructures localized to the distal end of a process were defined as terminal, while microstructures protruding from any other location along the length of the process were defined as mid-process. Distance from the cell body was measured using the Segmented Line tool in Fiji and defined as the shortest continuous path from the edge of the cell body to the most distal point of actin localization at the tips of filopodia, or most distal portion of actin protrusion for lamellipodia.

Filopodia and lamellipodia microstructures were further categorized by the presence or absence of GFAP within the microstructure. GFAP-positivity was defined by GFAP localization that extends to the actin-positive edge of the structure. Webbing-like microstructures were defined as filaments connecting two processes together and forming an enclosed extracellular space. For these, the distance from the cell body was defined using the Straight Line tool in Fiji from the edge of the cell body to the most distal webbing-like microstructure.

### Sample preparation for cryo-ET

R2/2 200 mesh quantifoil gold grids (Electron Microscopy Sciences) were cleaned with chloroform, placed in a sterile 35-mm glass-bottom dish (MatTek Life Sciences), glow-discharged using a Tergeo-EM plasma cleaner (PIE Scientific), and coated with poly-D-Lysine overnight. The dishes were washed three times with water before plating the astrocytes (20000 cells/dish) isolated using the procedure described above. The plated cells were incubated on the grid at 37 °C with 10% CO_2_ for 13 days as shown in Fig 2H. For cryo-ET study, the grid was blotted for 10 s with a blot force of 10 in a 95% humidity chamber of a Vitrobot Mark IV (Thermo Fisher Scientific) and vitrified by plunging into liquid ethane.

### Cryo-ET data acquisition

Data was acquired at the Cal-Cryo Facility at UC Berkeley using a Titan Krios G3i with a BIO Quantum energy filter (Gatan) operated at 300 kV (Thermo Fisher Scientific), using a K3 direct electron detector (Gatan). Tilt series were collected using Serial EM software (Mastronarde, 2005) at a magnification of 33,000x (pixel size of 2.63 Å), between -60° to + 60° tilt angles with 3° increments, an electron dose of 2.42 e^-^/Å^2^ per image (total dose of 99.62 e^-^/Å^2^), and at defocus values ranging between -3 to -6 µm.

### Cryo-ET data preprocessing and tomographic reconstruction

131 tilt series composed of 41 images each were acquired by manually selecting regions on primary processes of the astrocytes (as shown in Figure 4B and 4C). Beam-induced motion correction was performed on raw tilt images using Relion5 implementation of the UCSF MotionCor2 program(Scheres, 2012; Zheng et al., 2017), and the x8 binned images were then aligned for 3D reconstructing using automatic alignment in the Eman2 software, followed by CTF (Contrast Transfer Function) estimation to screen for good quality tomograms (Chen et al., 2019). Fifty-five selected tomograms were reconstructed at bin x4 (10.52 Å/pixel) using the Batch Tomogram Reconstruction, with Batchruntomo, and CTF-corrected using the Ctfplotter on IMOD (version 5.1)(Mastronarde, 2024; Mastronarde and Held, 2017). The reconstructed bin x4 tomograms were CTF deconvolved and corrected with IsoNet to enhance resolution and CTF correction, which help determine the microtubule polarity based on the protofilament skew(Liu et al., 2022).

### Tomogram segmentation

The cytoskeletal components of astrocytes in our tomograms were segmented using the non-commercial version of Dragonfly 2024.1 (Object Research Systems) following a previously described protocol (Heebner et al., 2022). Briefly, a gaussian filter was applied to the x4 binned tomogram to reduce noise and enhance signal. A small subregion containing cytoskeletal features was selected for manual annotation of about 7-10 tomographic slices. The segmentation wizard was then used to carry out iterative training of a 2.5-D U-Net segmentation model with 5 slices, initial filter count of 64, and depth level 5, until a good training model was obtained. This model was then used to predict the cytoskeletal features in whole tomograms. The predicted segmentations were manually cleaned up to eliminate segmented noise.

### Subtomogram averaging of microtubules

Microtubules from 18 different primary processes across 9 cells were used for averaging. From 31 x4 binned tomograms of primary processes, individual microtubules were manually traced by placing a marker at the approximate center using the *Volume Tracer* tool, and 3D particle picking was performed using the *Pick Particle* tool in Chimera (version 1.19)(Pettersen et al., 2004). Particle extraction and averaging was performed using subTOM (version 1.1.6) (https://github.com/DustinMorado/subTOM). The particles were extracted with a box size of 64 pixels centered on the coordinates picked on Chimera. Iterative alignment of the extracted subtomograms was carried out, followed by averaging. Z-slices of these averages were used to determine the microtubule protofilament number and the skew direction of the tubulin subunits that defines the polarity of the microtubule (Sosa and Chrétien, 1998; Chrétien et al., 1996). Of the 81 microtubule averages, 8 were inconclusive concerning protofilament number and/or polarity. The microtubule particle picking direction is key to determine the polarity: when the particle picking was from minus to the plus end, the averages show minus end polarity (clockwise direction). The protofilament skews of individual microtubules observed on the z-slices of the deconvolved tomograms therefore defined the microtubule polarity relative to the cellular compartment.

### Determination of MIP and lattice defect frequencies of individual microtubules

Microtubules from 12 different primary processes across 7 cells were used to determine MIP and lattice defect frequencies. The lengths of individual microtubules were measured on 21 selected good quality x8 binned tomograms of primary processes in Fiji. Good quality tomograms had the following characteristics: thin enough ice, minimal contamination, minimal distortions, good alignment, relatively high signal-to-noise ratio, and preservation of structural details. The number of MIPs and lattice defects in each microtubule were manually counted in 3dmod (version 5.1) using the slicer and the ZAP window (Mastronarde and Held, 2017). For each microtubule, the MIP and lattice defect frequencies were calculated by dividing their number by the microtubule length. The distance from the cell body to the primary process where the tomogram was acquired was estimated using Fiji in low resolution images of the area. Microtubules from tomograms of all the primary processes were pooled and categorized based on the distance from the cell body.

### Determination of filament abundance and lengths in astrocyte primary processes

Tomograms acquired on 10 different primary processes across 4 cells were used in this analysis. 11 good quality tomograms were used to analyze lengths and abundance of different classes of filaments. Of the 11 tomograms, 8 tomograms were used to generate data 10 – 20 µm, 5 at 0–10 µm, 2 at 20–30 µm, and 1 at 30– 40 µm from the cell body. Like microtubules, lengths of individual IF and actin filaments in tomograms acquired at the primary processes were measured in Fiji. Frequency counts of each filament type were used to generate a percentage abundance of each filament class in the region of tomogram acquisition.

### Statistics

Means, standard deviations, standard errors, and statistical analyses of immunofluorescent data were calculated and graphed using GraphPad Prism 10 (Version 10.4.0 (527)). Data were tested for normality or parametricity using Shapiro-Wilk and Kolmogorov-Smirnov tests. Normal data were analyzed using t-tests or ANOVAs while non-normal data were analyzed with Mann-Whitney or Kruskal-Wallis tests. In ANOVA analyses, repeated measures were used since multiple measurements from cells cultured together cannot be considered completely independent from one another. Box plots for both MIP and lattice defect frequencies were plotted and statistical pairwise tests were performed in R (R Core Team. 2025).

Violin plots represent median (dashed line) and 25^th^ and 75^th^ quartiles (dotted lines). Dots represent individual n’s, which are stated in the figure legends.

#### Description of supplementary material

Fig. S1 shows comparison of tubulin-based and GFAP-based primary branch numbers and lengths in primary immunopanned astrocytes. Fig. S2 shows the distribution of data for EB3-based estimates of run speed, duration, and length and more cryo-ET data on microtubule lattice defects. Fig. S3 shows the distribution of filament lengths and ratios of different filament types to one another, based on cryo-ET, as well as immunofluorescence data comparing the localization of GFAP and vimentin in immunopanned astrocytes. Fig. S4 provides examples of intermediate filaments and tubulin co-localizing by cryo-ET and extends the analysis of enrichment of GFAP relative to tubulin at branch tips that is shown in Fig. 6. Fig. S5 provides further image examples of actin webbing and extended analysis of the composition of actin microstructures in immunopanned astrocytes.

### Data availability

All data are available in the published article and its online supplemental material. Raw images or data underlying our statistical analyses are available from the corresponding author upon reasonable request.

## Supplementary Figures

**Fig. S1.**
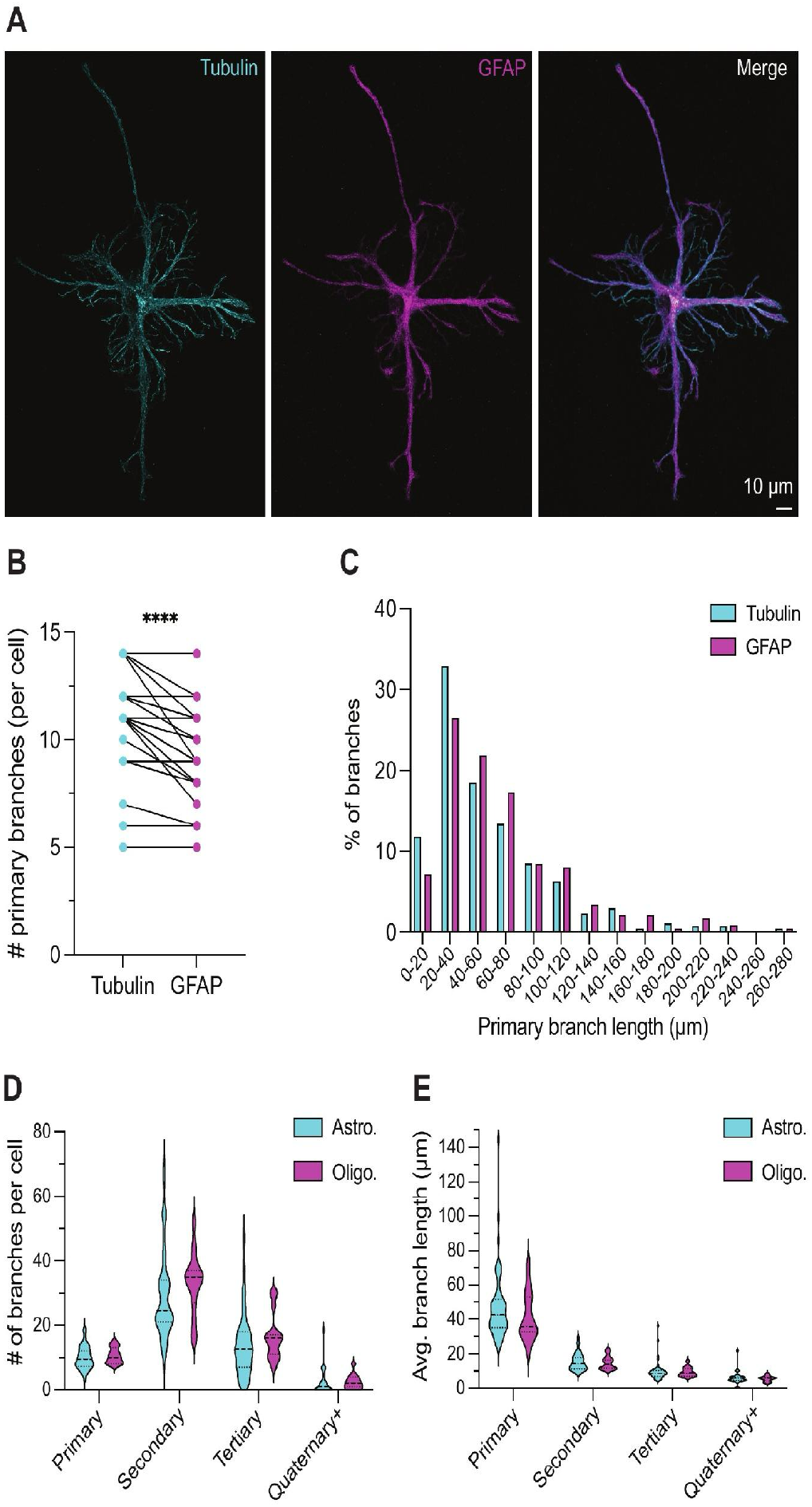
GFAP and tubulin label overlapping but distinct features of immunopanned astrocyte branch architecture. **(A)** Representative immunofluorescence images of a DIV14 immunopanned astrocyte stained for tubulin and GFAP. Scale bar, 10 µm. (B) Paired quantification of primary branch number per cell using tubulin or GFAP labeling. Each pair of connected points represents measurements from the same astrocyte. GFAP labeling detected fewer primary branches per cell than tubulin labeling. n = 25 cells from 3 independent cultures. Wilcoxon matched-pairs signed rank test, ****p ≤ 0.0001. **(C)** Frequency distribution of primary branch lengths measured plotted in (B) from tubulin- or GFAP-labeled branches, highlighting shift of distribution toward shorter lengths for tubulin. n = 270 total tubulin-labeled branches and 238 total GFAP-labeled branches. (D) Quantification of branch number per cell by branch order in immunopanned astrocytes and oligodendrocytes. Violin plots show median and interquartile range. (E) Quantification of average branch length by branch order in immunopanned astrocytes and oligodendrocytes. Average branch length was calculated per cell for each branch order present; cells lacking a given branch order were excluded from that branch-order length measurement. Data are from n = 44 astrocytes and n = 15 oligodendrocytes from 3-4 independent cultures/litters. Astrocyte data in (D, E) was replotted from Fig. 2C-D to show comparison with oligodendrocytes. Mixed-effects models with Tukey’s multiple comparisons testing showed no significant differences between cell types for branch order frequency or length.

**Fig. S2.**
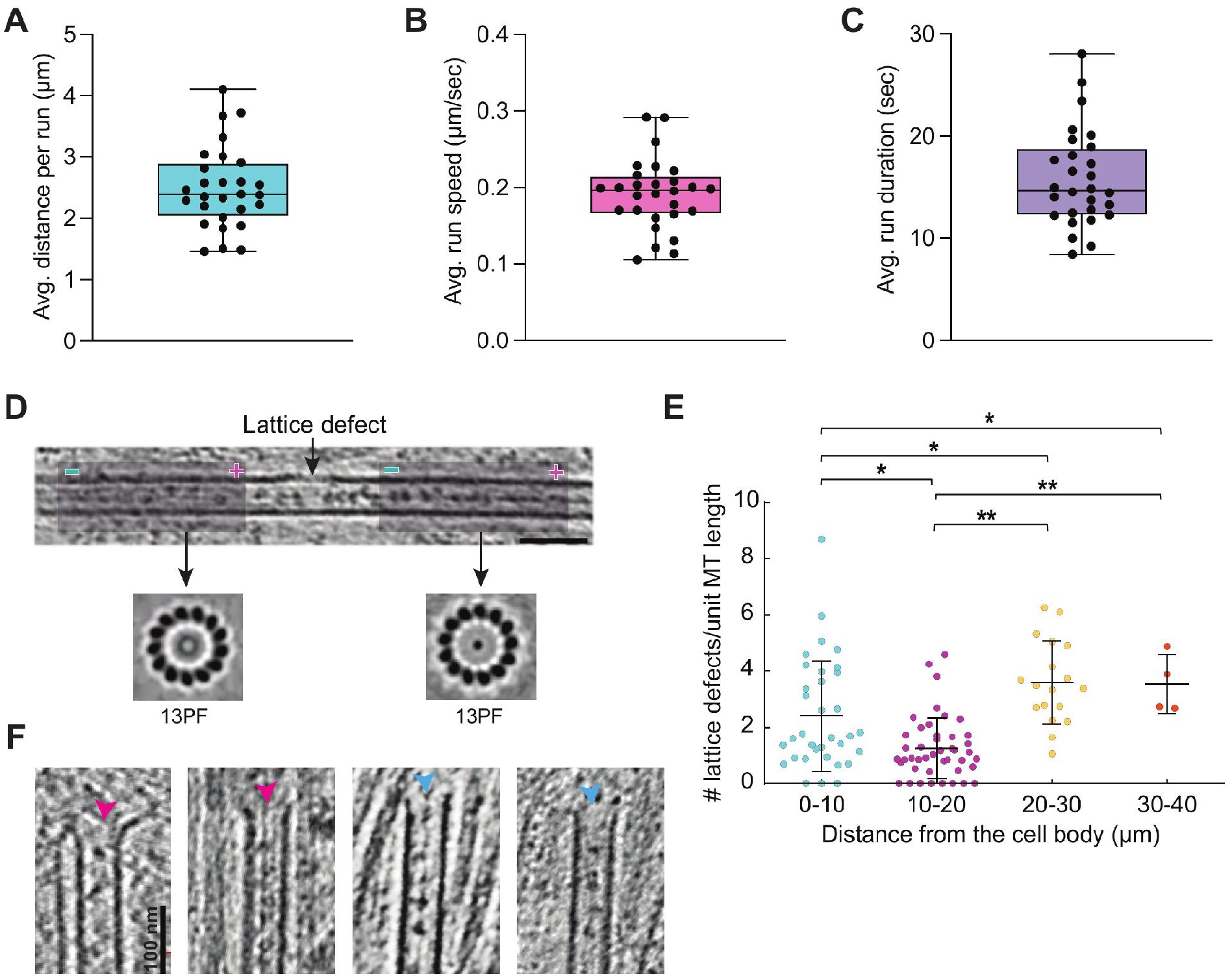
Extended analysis of astrocyte microtubules. **(A–C)** Box and whisker plots summarizing means per cell for run distance **(A)**, run speed **(B)**, and run duration **(C)** for EB3-mNeonGreen (EB3-mNG) comets in DIV13-DIV14 primary astrocytes expressing EB3-mNG. n = 110 kymographs from 28 cells from 3 independent cultures (3 litters of rats). **(D)** Tomogram slice containing a microtubule with a lattice defect (at the arrow, which points to missing protofilaments, as indicated by the lower gray level at the edge of the projection image). Below: end-on views of subtomogram averages showing maintenance of 13-protofilament geometry and plus-end-out polarity before and after the break. Dark purple boxes correspond to the areas used for averaging. Only microtubules with consistent averages along the lattice were analyzed. Scale bar: 100 nm. **(E)** Graph depicting the lattice defect frequency per µm microtubule length in astrocytic primary processes, binned by distance from the cell body: 0–10 µm (n = 35 microtubules), 10–20 µm (n = 46), 20–30 µm (n = 18), and 30–40 µm (n = 4). Bars represent mean ± SD. One-way ANOVA with Tukey’s multiple comparisons, ** p < 0.01. **(F)** Representative tomogram examples of microtubule ends highlighting slightly flared (pink arrowheads) and blunt (blue arrowheads) ends. Scale bars: 100 nm.

**Fig. S3.**
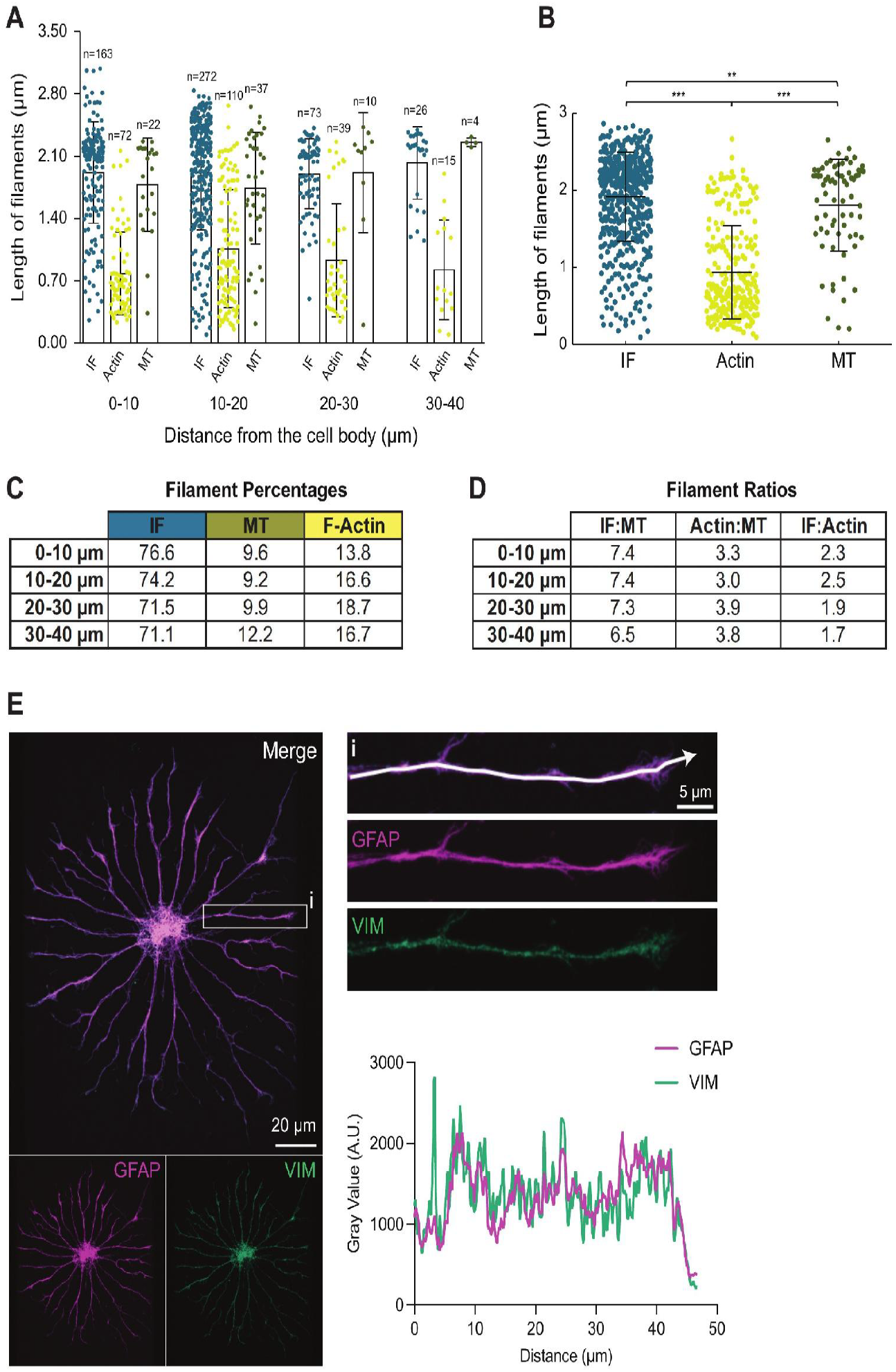
Composition of filament classes in astrocyte primary processes and similar localization patterns of GFAP and vimentin. **(A)** Bar graph showing the lengths of different filament classes measured in tomograms acquired in DIV13 astrocytic primary processes, binned by distance from the cell body: 0–10 µm, 10–20 µm, 20–30 µm, and 30–40 µm. n above each bar shows the number of individual filaments analyzed. **(B)** Bar graph showing average length of different filament classes in primary astrocyte processes, pooling all filaments together analyzed in (A). One-way ANOVA with Tukey’s multiple comparisons. Bars in (A) and (B) represent mean ± SD. **(C)** Table depicting percentage abundance of each filament class in primary astrocyte processes, calculated with raw numbers of filaments in (A) and binned by distance from the cell body: 0–10 µm, 10–20 µm, 20–30 µm, and 30–40 µm. **(D)** Table showing ratios of different filament classes to one another in primary astrocyte processes, calculated with raw numbers of filaments in (A) and binned by distance from the cell body: 0–10 µm, 10–20 µm, 20–30 µm, and 30–40 µm. **(E)** Representative immunofluorescence staining image of GFAP and vimentin in DIV13 primary astrocytes, showing overlapping localization of these IFs. Inset highlights one branch with a line scan quantification of fluorescent intensity value for the branch depicted below, showing similar intensity patterns along the branch length for GFAP and vimentin.

**Fig. S4.**
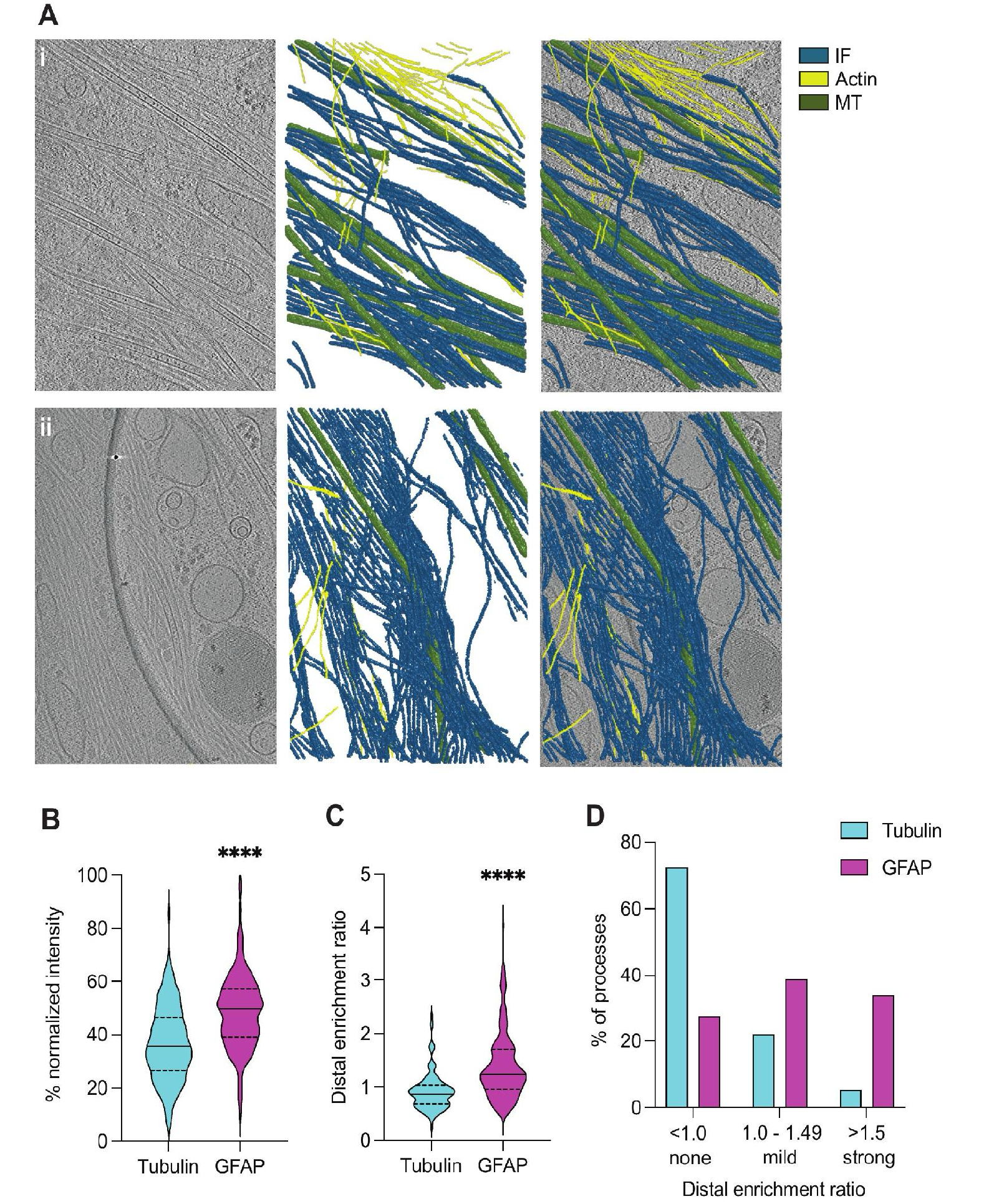
Interplay between microtubules and IFs observed by cryo-ET and extended analysis of distal GFAP enrichment in astrocyte process tips. **(A)** 2D slices of two example tomograms (i, ii) highlighting the presence of IFs alongside microtubules. In both i and ii, the rightmost image shows a 2D slice of a reconstructed and CTF-corrected tomogram, the middle image shows segmentation of cytoskeletal filaments based on neural-network training, and the left image shows segmentation overlaid on the tomogram. Scale bar: 50 nm. **(B)** Violin plot depicting the percent intensity of tubulin and GFAP across the distal 10 µm of primary and secondary astrocyte processes. Intensities were normalized against the maximum value across the entire branch. Mean ± SEM for tubulin and GFAP, respectively, are 36.8 ± 1.2% and 49.6% ± 1.1%. **(C)** Violin plots showing significantly more distal enrichment of GFAP compared to tubulin. Distal enrichment ratio was calculated as the average % normalized intensity in the distal 10 µm of process tips divided by the average % normalized intensity in the preceding 40 µm. Mean ± SEM for tubulin and GFAP, respectively, are 0.91 ± 0.03 and 1.40 ± 0.05. **(D)** Bar graph showing percentage of processes classified by variable distal enrichment ratios. Enrichment ratios below 1.0 correspond to lower intensity in the distal 10 µm. Mild enrichment ratios correspond to <50% higher intensity across the distal 10 µm. Strong distal enrichment ratios correspond to >50% higher intensity in the distal 10 µm. (B) and (C) analyzed with Mann-Whitney test, **** p ≤ 0.0001. For (B–D), n = 141 processes from 25 cells across 3 independent experiments (from 3 litters of rats).

**Fig. S5.**
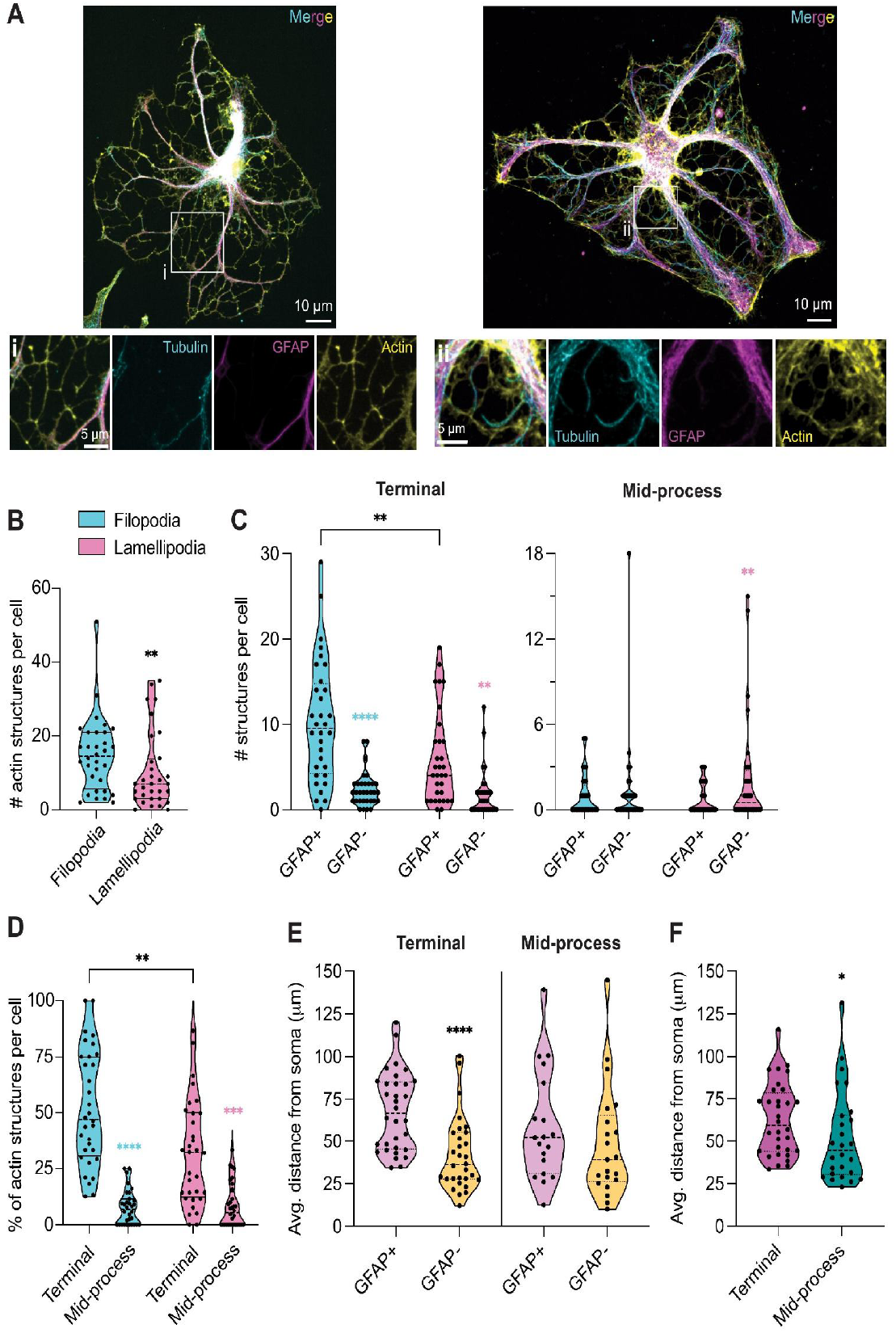
Expanded analysis of actin microstructures and GFAP co-localization in astrocyte processes. Violin plots comparing: (A) Additional immunofluorescence images showing examples of actin-based reticular webbing. Immunopanned astrocytes (DIV13-DIV14) were stained for tubulin, GFAP, and F-actin using phalloidin. Actin forms reticular web-like structures surrounding the soma and extending between major astrocyte branches. Boxed regions are enlarged below and show that reticular webbing is strongly labeled by phalloidin and often contained less GFAP and tubulin signal than the thicker branch shafts. Scale bars, 10 µm in whole-cell images and 5 µm in magnified insets. Violin plots comparing: **(B)** Total numbers of filopodia and lamellipodia per astrocyte. **(C)** Numbers of GFAP-positive versus GFAP-negative filopodia or lamellipodia per astrocyte, segregated by location. In the terminal, GFAP-positive filopodia (mean ± SEM: 10.3 ± 1.3 per cell) significantly outnumbered GFAP-positive lamellipodia (5.8 ± 1.0); no difference was observed between the numbers of GFAP-negative filopodia (2.3 ± 0.4) and GFAP-negative lamellipodia (1.8 ± 0.5). At mid-process sites, lamellipodia were more often GFAP-negative (2.1 ± 0.7) than GFAP-positive (0.7 ± 0.2). **(D)** Average percentages per cell of different types of actin structures observed, segregated by filopodia vs. lamellipodia along with terminal vs. mid-process location. For both filopodia and lamellipodia, terminal structures were significantly more abundant (Terminal filopodia: 52.1 ± 4.53%, mid-process filopodia: 7.7 ± 1.36%, terminal lamellipodia: 31.9 ± 4.13%, and mid-process lamellipodia: 8.3 ± 1.73%; mean ± SEM) **(E)** Average distance per astrocyte from the edge of the soma to the actin microstructure segregated by terminal versus mid-process location and the presence or absence of GFAP **(F)** Average distance per astrocyte from the edge of the soma to the actin microstructure, segregated only by terminal versus mid-process location. Mann-Whitney test or 2-way ANOVA with repeated measures followed by Tukey’s multiple comparisons testing. n = 33 cells from 4 independent cultures (from 4 litters of rats). * p ≤ 0.05, ** p ≤ 0.01, *** p ≤ 0.001, **** p ≤ 0.0001. In (C) and (D), cyan asterisks are used for within filopodia comparisons and pink asterisks are used for within lamellipodia comparisons.

### Supplementary Movie

**Movie S1**. Example live movie of astrocytes transfected with EB3-mNeonGreen to label growing microtubule plus ends. Movie was acquired using 3 second intervals over 3 minutes total. Framerate is 10 FPS.

